# A phage nucleus-associated RNA-binding protein is required for jumbo phage infection

**DOI:** 10.1101/2023.09.22.559000

**Authors:** Eray Enustun, Emily G. Armbruster, Jina Lee, Sitao Zhang, Brian A. Yee, Yajie Gu, Amar Deep, Jack T. Naritomi, Qishan Liang, Stefan Aigner, Benjamin A. Adler, Brady F. Cress, Jennifer A. Doudna, Vorrapon Chaikeeratisak, Don W. Cleveland, Majid Ghassemian, Gene W. Yeo, Joe Pogliano, Kevin D. Corbett

## Abstract

Large-genome bacteriophages (jumbo phages) of the *Chimalliviridae* family assemble a nucleus-like compartment bounded by a protein shell that protects the replicating phage genome from host-encoded restriction enzymes and CRISPR/Cas nucleases. While the nuclear shell provides broad protection against host nucleases, it necessitates transport of mRNA out of the nucleus-like compartment for translation by host ribosomes, and transport of specific proteins into the nucleus-like compartment to support DNA replication and mRNA transcription. Here we identify a conserved phage nuclear shell-associated protein that we term Chimallin C (ChmC), which adopts a nucleic acid-binding fold, binds RNA with high affinity *in vitro*, and binds phage mRNAs in infected cells. ChmC also forms phase-separated condensates with RNA *in vitro*. Targeted knockdown of ChmC using mRNA-targeting dCas13d halts infections at an early stage. Taken together, our data suggest that the conserved ChmC protein acts as a chaperone for phage mRNAs, potentially stabilizing these mRNAs and driving their translocation through the nuclear shell to promote translation and infection progression.

## Introduction

The continual arms race between bacteria and bacteriophages (phages) has driven the development of myriad immune systems in bacteria, along with an equally complex set of phage-encoded immune countermeasures (1, 2). A striking example of these immune countermeasures is the nucleus-like compartment assembled by a family of large-genome “jumbo” phages (defined as phages with genomes >200 kb in size) now termed *Chimalliviridae* (3–7). This compartment shields the phages’ replicating genomes from host-encoded defenses including restriction enzymes and CRISPR/Cas nucleases (8, 9) . The phage nuclear boundary or shell primarily comprises a single protein, termed Chimallin (ChmA), which assembles into a lattice with pores less than ∼2 nm in size, which can allow the passage of metabolites but restrict the passage of most proteins (7, 10). Most *Chimalliviridae* also encode a tubulin homolog called PhuZ, which assembles into dynamic filaments that center and rotate the phage nucleus within the infected cell while also trafficking pro-capsids to the phage nucleus for genome packaging (3–5).

The physical barrier erected by ChmA-encoding jumbo phages between their replicating genomes and the host cytoplasm introduces a number of challenges to the phage that mirror challenges faced by eukaryotic cells and their nuclei. In particular, since mRNAs are produced within the phage nucleus but the translation machinery is located in the host cytoplasm, mRNAs must be translocated out of the phage nucleus for translation. Similarly, any phage protein whose function requires it to be localized within the phage nucleus must be specifically translocated from the cytoplasm into the phage nucleus after translation. Finally, replicated phage genomic DNA must be translocated through the nuclear shell for packaging into pro-capsids that are docked onto the exterior of the nuclear shell (4, 5, 11, 12).

The requirement for translocation of mRNAs, proteins, and genomic DNA through the phage nuclear shell implies the existence of additional shell components embedded within or associated with the ChmA lattice that mediate these activities. In prior work, we used proximity labeling and localization analysis in the *Chimalliviridae* family phage PhiPA3 to identify proteins that physically associate with ChmA and localize to the nuclear shell (13). One of these proteins, termed ChmB, interacts directly with ChmA both *in vitro* and *in vivo*, and associates with the virion portal protein *in vitro*. ChmB’s network of protein-protein interactions and its distinctive 3D structure suggest that it may form pores in the phage nuclear shell that enable the docking of pro-capsids for genome packaging. Further data showing that the expression of dominant-negative ChmB mutants compromises early steps in phage nucleus growth and maturation further suggests that ChmB may participate in mRNA and/or protein translocation through the nuclear shell (13).

Here, we show that another conserved phage nuclear shell-associated protein, which we term ChmC, adopts a nucleic acid-binding fold and binds RNA *in vitro*, and forms RNA-protein condensates through a conserved asparagine-rich C-terminal region. In phage-infected cells, ChmC specifically binds phage mRNAs. Targeting ChmC using Cas13-based translational knockdown reveals that the protein plays critical roles in assembly, growth, and maturation of the phage nucleus. Together, these data suggest that ChmC acts as a chaperone for phage-encoded mRNAs, likely aiding their translocation through the nuclear shell to promote translation and infection progression.

## Results

### Identification of the abundant and early-expressed jumbo phage protein ChmC

We previously used proximity ligation to identify proteins in the nucleus-forming jumbo phage PhiPA3 that physically associate with the major nuclear shell protein ChmA or with a phage nucleus-localized protein, UvsX (gp175) (13). Through this analysis, we identified a minor nuclear shell component (ChmB) and several uncharacterized proteins with no known function (13). One of these proteins was PhiPA3 gp61, which is located in a highly conserved cluster of genes that includes *chmA* (gp53), several subunits of the phage-encoded “non-virion RNA polymerase” (nvRNAP: gp62, gp65-66, and gp67), and two additional phage nucleus-associated proteins, gp63 and gp64 (Figure 1A). gp61 is of particular interest as its homolog is the second highest-expressed non-structural protein in *Pseudomonas chlororaphis* cells infected with the related jumbo phage 201Phi2-1 (gp123) (4). To test whether PhiPA3 gp61 is also highly expressed, we infected *P. aeruginosa* cells with PhiPA3 and performed mass spectrometry proteomics to identify the timing and expression of phage proteins. We confirmed that ChmA is the most abundantly expressed non-structural protein, and that gp61 is the second-most abundant (Figure 1B, Table S1-S2). Based on its conservation, abundance, and association with the phage nuclear shell, we term this protein Chimallin C (ChmC).

**Figure 1.**
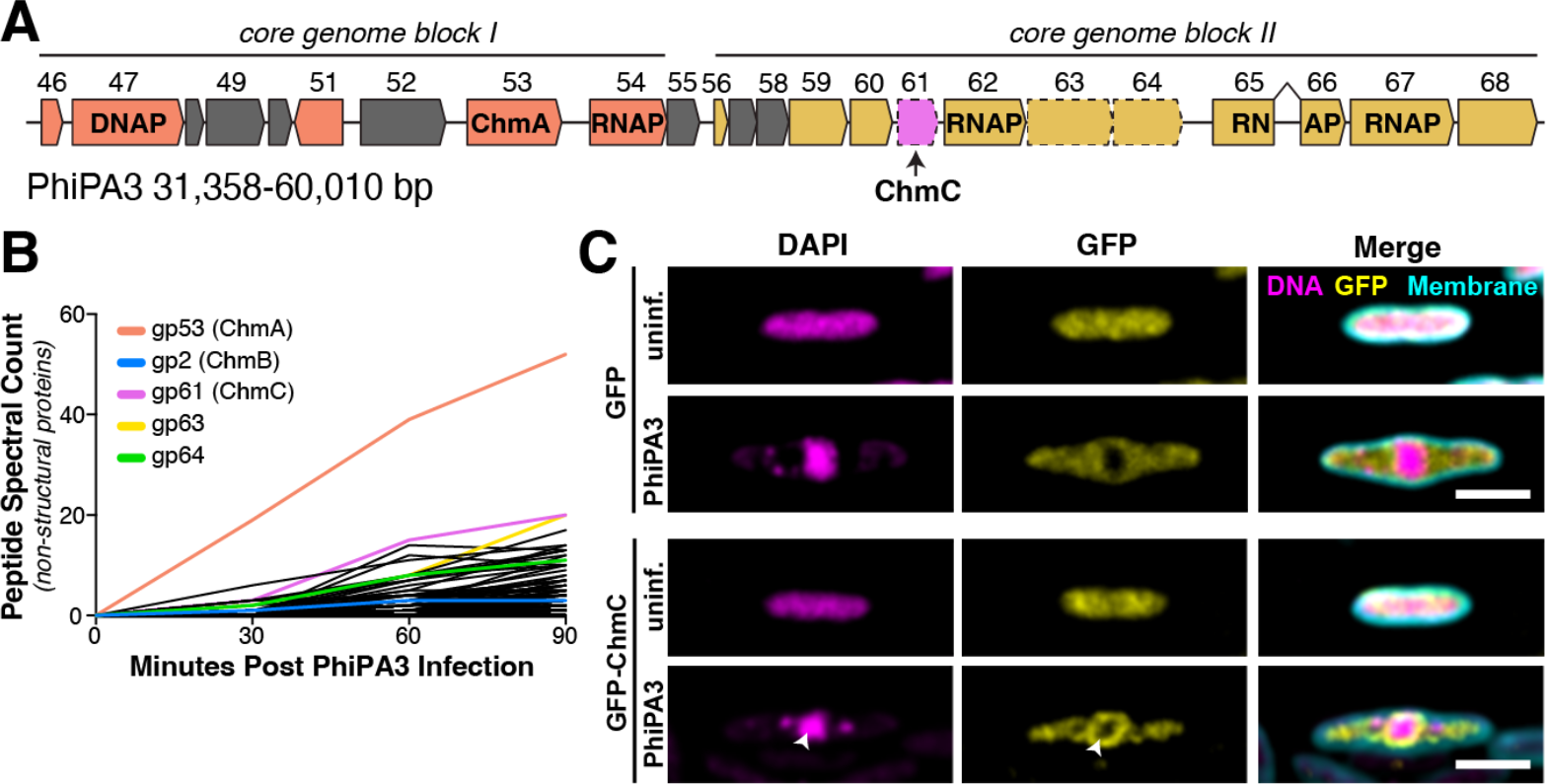
PhiPA3 gp61 is associated with the phage nucleus. **(A)** Map of the PhiPA3 genome spanning jumbo phage conserved blocks I and II (3). Genes in conserved block I are colored salmon, and genes in conserved block II are colored goldenrod. Genes conserved with the closely related jumbo phage PhiKZ but not all jumbo phages are colored gray, and non-conserved genes are colored white. gp47 is a putative DNA polymerase (DNAP), and gp53 is the major nuclear shell protein Chimallin (ChmA). Gp54, gp64, gp65-66 (interrupted by a self-splicing intron), and gp67 are subunits of the phage-encoded non-virion RNA polymerase (RNAP). Dotted outlines indicate three proteins (gp61, gp63, and gp64) shown to be associated with the phage nuclear shell (13). **(B)** Mass spectrometry proteomics analysis of PhiPA3-infected *P. aeruginosa*, showing spectral counts of non-structural phage proteins. ChmA (gp53), ChmB (gp2), gp61, gp63, and gp64 are shown in colors and labeled. See **Table S1-S2** for full mass spectrometry results. **(C)** Localization of sfGFP (top) or sfGFP-fused PhiPA3 gp61 (bottom) in PhiPA3-infected *P. aeruginosa* cells at 75 minutes post infection. Magenta: DAPI nucleic acid dye; cyan: FM4-64 membrane dye; yellow: GFP. Arrowheads indicate phage nuclei, and asterisks indicate phage bouquets. Scale bar = 2 μm.

Confirming our earlier microscopic observations (13), we find that ectopically-expressed PhiPA3 ChmC fused to GFP localizes to the phage nuclear shell in late-stage PhiPA3 infections of *P. aeruginosa* (Figure 1C). Fluorescent microscopy is unable to determine whether ChmC is localized within the nucleus or just outside the ChmA shell, but our prior identification of ChmC through proximity-labeling with the nuclear-localized protein UvsX suggests that the protein is at least partially localized within the phage nucleus (13). We next purified GFP-tagged ChmC from *P. aeruginosa* cells infected with PhiPA3 and used mass spectrometry to identify associated proteins (Figure S1, Table S3). In this analysis, we identified the putative phage nucleus pore protein ChmB (gp2) and one subunit of each phage-encoded RNA polymerase: gp62 is part of the non-virion RNA polymerase, and gp77 is part of the virion RNA polymerase. We also observed association with 19 ribosomal and ribosome-associated proteins plus the host RNA polymerase α, β, and β′ subunits. The enrichment of ribosomal proteins and RNA polymerase subunits in this experiment suggests that ChmC may be a non-specific RNA binding protein. The observed association of ChmC with the putative pore-forming protein ChmB, moreover, suggests that these two proteins may functionally cooperate at the nuclear shell.

### ChmC adopts a nucleic acid-binding fold and binds RNA

ChmC is conserved across jumbo phages but shows no identifiable sequence similarity to known proteins. To gain insight into ChmC’s structure and potential function, we used AlphaFold2 (14) to predict its 3D structure with high confidence (Figure 2A-B, Figure S2A-B). Analysis of the resulting model using the DALI protein structure comparison server revealed a predicted Whirly domain fold (also termed a PUR domain) common to multiple families of single-stranded RNA and DNA binding proteins(15–17). The Whirly fold typically comprises a tandem repeat of β-β-β-β-α secondary structure elements, and is exemplified by the *Trypanosoma brucei* MRP1 protein (Figure 2A) (16). The predicted structure of PhiPA3 ChmC shows a tandem repeat of β-β-β-α elements, with an overall 3D structure highly reminiscent of the Whirly domain (Cα r.m.s.d. of 4.7 Å for gp61 versus *T. brucei* MRP1 over 120 residues) (Figure 2A). Whirly domain proteins typically form higher-order complexes including homo- and heterotetramers, exemplified by the 2:2 heterotetramers formed by *T. brucei* MRP1 and MRP2 (16). We used AlphaFold 2 to predict the structure of PhiPA3 ChmC oligomers, and obtained a confident prediction of a ChmC homotetramer that is strikingly similar to the *T. brucei* MRP1:MRP2 heterotetramer structure (Figure 2B, Figure S2A, C). We expressed and purified full-length PhiPA3 ChmC in *E. coli* and analyzed its oligomeric state by size exclusion chromatography coupled to multi-angle light scattering (SEC-MALS). Supporting our structure prediction, we found that ChmC forms a stable homo-tetramer in solution (Figure 2C).

**Figure 2.**
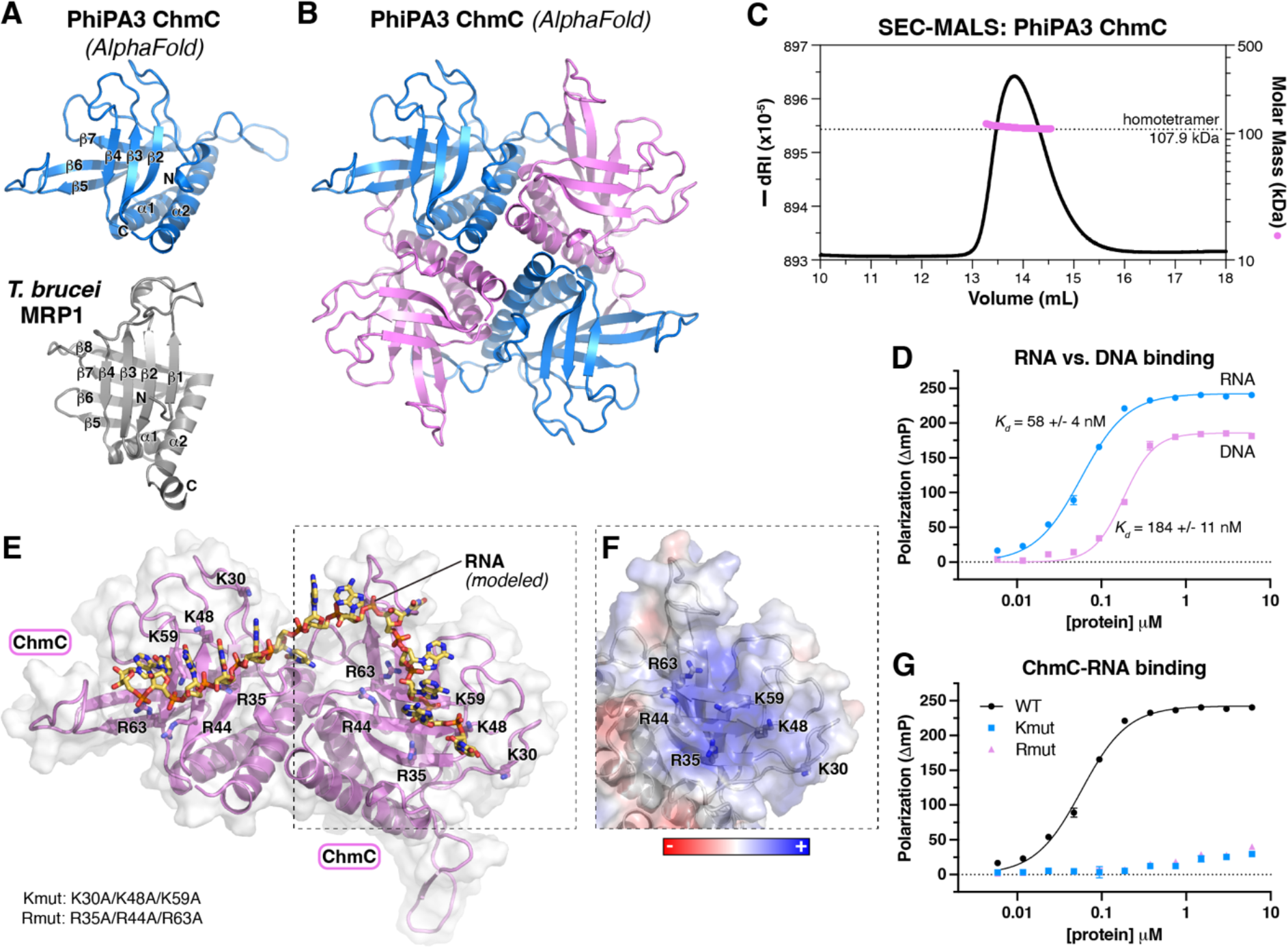
ChmC adopts an RNA-binding Whirly fold. **(A)** AlphaFold predicted structure of PhiPA3 ChmC (gp61; blue) at top oriented equivalently to the structure of *T. brucei* MRP1 (PDB ID 2GJE; gray) (16) at bottom, with secondary structure elements labeled. **(B)** AlphaFold predicted structure of a PhiPA3 ChmC homotetramer, with alternating subunits colored blue and pink. See **Figure S2A-B** for details of AlphaFold predictions. **(C)** Size exclusion chromatography coupled to multi-angle light scattering (SEC-MALS) analysis of purified PhiPA3 ChmC. **(D)** Single-stranded RNA (blue circles) and DNA (pink squares) binding of PhiPA3 ChmC as measured by fluorescence polarization. Data points are shown as average +/-standard deviation of triplicate technical replicates, and curves are fit with a cooperative binding model (Hill coefficient for RNA binding 1.6 +/-0.1; Hill coefficient for DNA binding 2.4 +/-0.3). **(E)** Structural model of PhiPA3 ChmC binding single-stranded RNA, generated by overlaying two adjacent subunits of the ChmC AlphaFold 2 model shown in panel (B) with structure of a *T. brucei* MRP1:MRP2 heterotetramer bound to RNA (PDB ID 2GJE; see **Figure S2C**) (16). Lysine and arginine residues mutated to generate the Kmut (K30A/K48A/K59A) and Rmut (R35A/R44A/R63A) are shown as sticks and labeled. **(F)** Surface charge representation of one ChmC subunit, oriented equivalently to panel (E) (dotted line). See **Figure S2D. (G)** Single-stranded RNA binding of PhiPA3 ChmC wild-type (black circles, reproduced from panel (D)), Kmut (blue squares), and Rmut (pink triangles), as measured by fluorescence polarization. Data points are shown as average +/-standard deviation of triplicate technical replicates. **See Figure S2E** for DNA binding.

Given the known roles of Whirly domain proteins in nucleic acid binding, we tested the ability of purified PhiPA3 ChmC to bind single-stranded DNA or RNA *in vitro*. Using a fluorescence polarization assay, we found that ChmC binds both DNA and RNA, but that the protein binds RNA with a higher affinity (*K*_*d*_ = 58 +/-4 nM) than it binds DNA (*K*_*d*_ = 184 +/-11 nM) (Figure 2D). We modeled a ChmC-RNA complex by overlaying the PhiPA3 ChmC tetramer model onto the structure of *T. brucei* MRP1:MRP2 bound to RNA(16) (Figure 2A, S2C). Based on this model, we designed two multi-site mutants to disrupt nucleic acid binding, termed Kmut (K30A/K48A/K59A) and Rmut (R35A/R44A/R63A) (Figure 2E-F, S2D), and found that both mutants completely disrupt RNA and DNA binding *in vitro* (Figure 2G, S2E). Based on these data, we conclude that gp61 adopts a homotetramer of Whirly domain folds and binds RNA.

### ChmC forms condensates with RNA

AlphaFold2 structure predictions of ChmC homologs from different phages consistently showed low confidence scores (pLDDT) for the C-terminal 50-70 residues of the protein, indicating that this region is likely disordered in solution. Across several ChmC orthologs, this region contains a ∼40-residue subregion enriched in asparagine and glycine residues (N/G-rich), followed by a ∼15-residue subregion enriched in serine, aspartate, and glutamate residues (Figure 3A). N/G-rich regions have been shown to promote macromolecular condensate formation, likely through dipole-dipole interactions (18, 19). Disordered regions rich in serine, aspartate, and glutamate, meanwhile, have been termed “electronegative clusters” (ENCs) which can stabilize RNA binding proteins in solution and suppress nonspecific RNA binding (20). Indeed, both the PSPredictor and catGRAN-ULE algorithms strongly predicted that ChmC forms condensates, and that this propensity is driven by the protein’s C-terminal disordered region (Figure S3A). Together with our finding that ChmC binds RNA *in vitro*, these predictions suggested that ChmC might form macromolecular condensates with RNA through multivalent RNA binding and low-affinity interactions through its C-terminal disordered region, similar to many other RNA binding proteins in both prokaryotes (21, 22) and eukaryotes (23–25). To directly test the propensity for PhiPA3 ChmC to form condensates, we engineered a cysteine residue at the protein’s N-terminus to enable fluorescent labeling using a maleimide-linked Cy5 dye. In a low-salt buffer (50 mM NaCl), we observed that purified ChmC forms uniform small droplets, and that the addition of a 2.3 kb RNA with the major capsid protein (gp136) gene sequence stimulates formation of large droplets that showed hallmarks of liquid-liquid phase separation, including dynamic growth and fusion of droplets (Figure 3B, S3B). Disrupting ChmC’s ability to bind RNA using the Kmut or Rmut multisite mutations dramatically reduced, but did not eliminate formation of condensates in the presence of RNA (Figure 3B). Since both the Kmut and Rmut proteins retain some positively charged residues on the RNA binding surface, it is likely that these proteins retain some ability to bind RNA, albeit with lower affinity than wild-type ChmC. Removal of the C-terminal disordered region (residues 204-251 removed; ChmC-ΔC), meanwhile, completely eliminated formation of condensates in both the absence and presence of RNA (Figure 3B). Importantly, the ChmC-ΔC protein formed homotetramers and retained the ability to bind RNA *in vitro* (Figure S3C-D).

**Figure 3.**
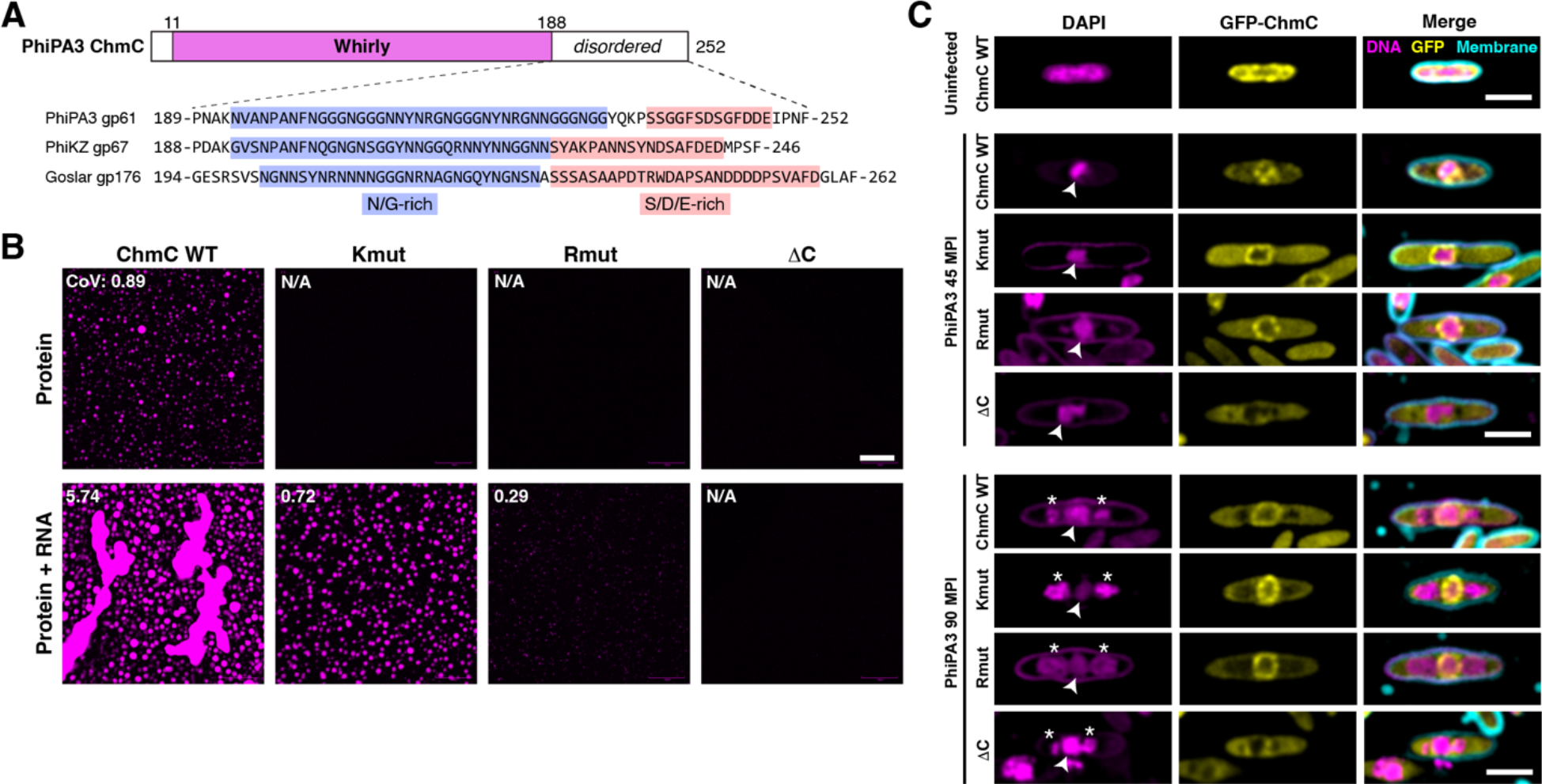
ChmC forms phase-separated condensates with RNA. **(A)** *Top:* Domain structure of PhiPA3 ChmC, with Whirly domain colored magenta and regions predicted to be disordered in white. Bottom: Sequence of the C-terminal predicted disordered domain in ChmC from PhiPA3 (gp61), PhiKZ (gp67), and Goslar (gp176). Asparagine/glycine (N/G) rich regions are highlighted in blue, and serine/aspartate/glutamate (S/D/E) rich regions are highlighted in salmon. See **Figure S3A** for catGRANULE analysis of all three proteins, and **Figure S3C-D** for analysis of ChmC ΔC binding DNA and RNA. **(B)** Fluorescence microscopy imaging of PhiPA3 ChmC (wild type, Kmut, Rmut, or ΔC; 10% Cy5-labeled) at 30 μM protein concentration, either alone (top row) or with 2 μM of a 40-base RNA (2.8 μg/mL; middle row) or 83 nM of a 2.3 kb RNA (5.8 μg/mL; bottom row). All images were taken 30 minutes after final dilution and mixing with RNA. For all conditions that showed condensate formation, the coefficient of variation (CoV) was calculated as the standard deviation of particle area divided by the mean particle area (WT protein alone n=423; WT + 40 base RNA n=222; Kmut + 40 base RNA n=247; Rmut + 40 base RNA n=5; WT + 2.3 kb RNA n=552; Kmut + 2.3 kb RNA n=368; Rmut + 2.3 kb RNA n=147). Scale bar = 30 μm. See **Figure S3B** for DIC imaging. **(C)** Localization of GFP-tagged PhiPA3 ChmC (wild type, Kmut, Rmut, or ΔC in PhiPA3-infected *P. aeruginosa* cells at 45 and 110 minutes post infection (MPI). Yellow: GFP; magenta: DAPI nucleic acid; cyan: FM4-64 membrane dye. Arrowheads indicate phage nuclei, and asterisks indicate phage bouquets. Scale bar = 2 μm.

We next examined localization of ChmC mutants in PhiPA3-infected *P. aeruginosa* cells. The RNA binding mutant proteins (Kmut and Rmut) showed localization to the nuclear shell in infected *P. aeruginosa* cells, equivalent to wild-type protein. A limitation of this assay is that the mutant proteins are expressed alongside wild-type phage-encoded ChmC, and may form hetero-oligomers with wild-type ChmC in infected cells. Despite this limitation, both the Kmut and Rmut showed a localization pattern distinct from wild-type ChmC, retaining nuclear shell localization but showing more uniform cytoplasmic distribution than wild-type protein (Figure 3C). Strikingly, removal of the ChmC C-terminus completely disrupted nuclear shell binding activity, suggesting that the protein’s ability to form condensates (with or without RNA) is important for the protein’s localization to the nuclear shell. Together with our *in vitro* RNA binding and RNA-mediated condensation data, these localization data suggest that ChmC forms RNA-protein condensates associated with the nuclear shell, and potentially also in the host cytoplasm.

### ChmC associates with viral mRNAs in infected cells

To test whether ChmC associates with RNA, particularly phage mRNAs, in infected cells, we performed eCLIP-seq (enhanced UV crosslinking and immunoprecipitation, followed by deep sequencing) in PhiPA3-infected *P. aeruginosa* cells expressing GFP-tagged ChmC. We obtained ∼1.6M uniquely-mapping sequence reads for GFP-tagged PhiPA3 ChmC across two independent replicates (1,161,658 + 423,427 reads), and 1.5M reads for control GFP samples (1,070,428 + 423,357 reads). We mapped all reads to the host (*P. aeruginosa* PA01) and phage genomes (Figure 4A, S4A), then calculated the enrichment for each gene for immuno-precipitated samples compared to matched input samples. All samples were collected 45 minutes after phage infection, after the host genome is fully degraded and the phage genome has begun replicating (4, 5, 11).

**Figure 4.**
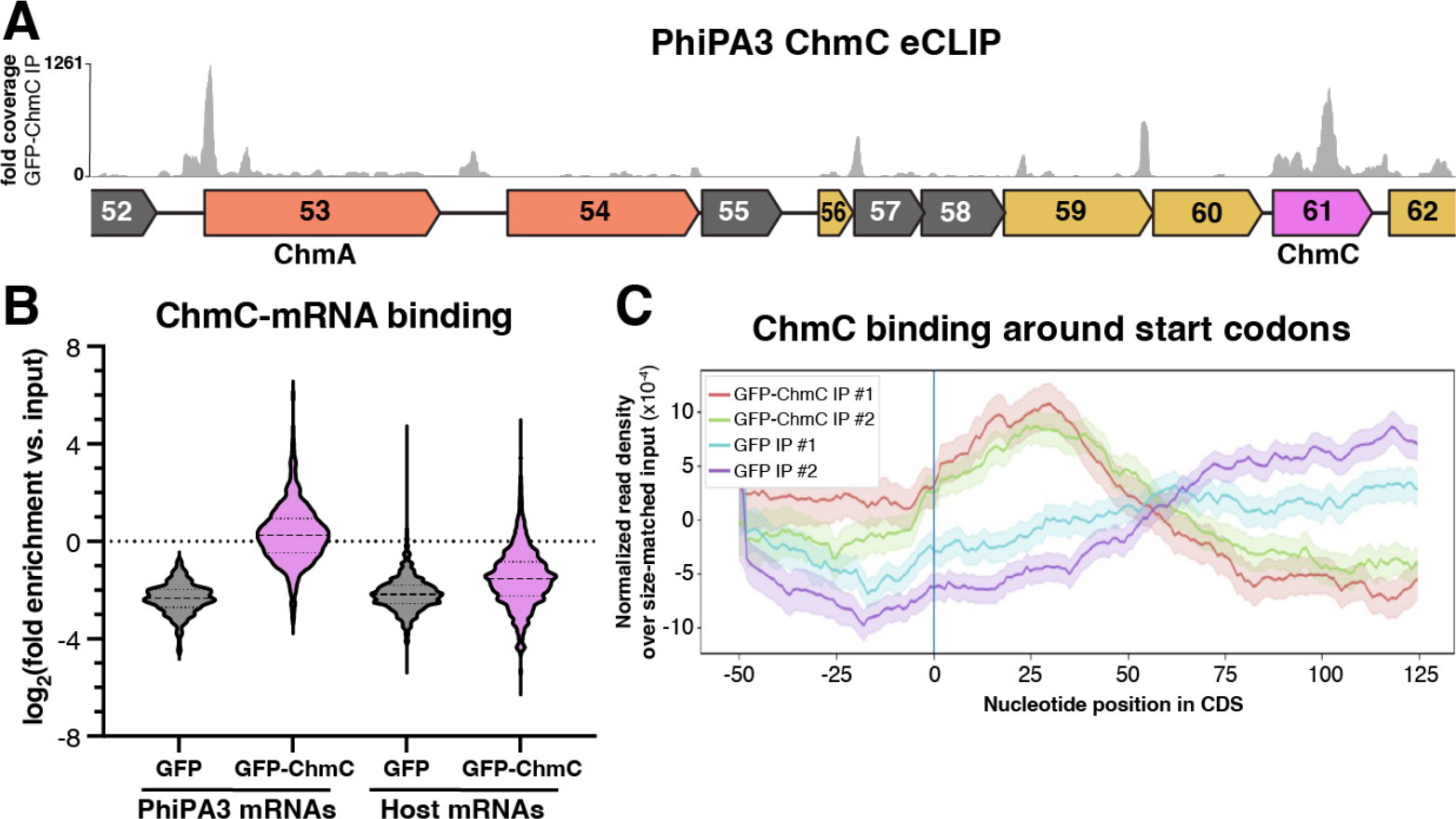
ChmC binds phage mRNAs. **(A)** Fold sequence coverage from eCLIP analysis of GFP-ChmC, in the region of the PhiPA3 genome encoding ChmA and ChmC. See **Figure S4** for eCLIP validation. **(B)** Log_2_(fold enrichment versus input for all PhiPA3 and host (P. aeruginosa) genes detected in eCLIP analysis of GFP (gray) and GFP-ChmC (pink). **(C)** Metagene analysis showing enrichment of GFP-ChmC binding near start codons of PhiPA3 genes (replicate #1 red, replicate #2 green).

*Chimalliviridae* encode two multi-subunit RNA polymerases: a “virion RNA polymerase” (vRNAP) that is packaged in the virion and is responsible for transcription immediately after infection, and a “non-virion RNA polymerase” (nvRNAP) that is responsible for transcription in the middle and late stages of infection(26–28). A prior RNA-seq analysis of the jumbo phage PhiKZ defined the operon structure of this phage and found that while vRNAP-transcribed early genes have a defined promoter sequence, genes transcribed by the nvRNAP do not show a reproducible promoter sequence (29). This analysis also identified a number of noncoding antisense RNAs, which were proposed to regulate translation of phage genes (29). Since a similar RNA-seq analysis has not been performed for PhiPA3, our analysis of gp61 eCLIP data is limited to annotated protein- and tRNA-encoding genes. We first analyzed the relative enrichment of phage mRNAs versus host mRNAs in the ChmC eCLIP immunoprecipitates versus input RNA. We found that phage mRNAs were slightly enriched (median log_*2*_(fold enrichment) of 0.186) compared to input samples, while host-encoded mRNAs were significantly depleted (median log_*2*_(fold enrichment) of -1.54) compared to input (Figure 4B). Overall, these data suggest that ChmC preferentially associates with phage mRNAs compared to host mRNAs, but that within phage mRNAs, ChmC shows little to no specificity. This preference may arise from ChmC’s localization within the phage nucleus, and/or from an inherent specificity for particular sequences or structures in phage mRNAs.

We next visually inspected ChmC eCLIP sequence coverage on the PhiPA3 genome. While mRNAs for some highly expressed genes (judging from the abundance of sequence reads in eCLIP input samples) were bound by ChmC across the entire open reading frame (e.g. the *chmC* gene itself; Figure 4A), we noticed that the majority of binding occurred in defined peaks near the start codons of genes. While the operon structure of PhiPA3 is not annotated, many peaks occurred near the start codons of genes that appear to be within polycistronic mRNAs. That is, ChmC binding occurs not only near the 5’ end of an mRNA (e.g. *chmA*; Figure 4A), but also likely occurs near internal start codons in mRNAs that encode multiple genes (e.g. gp57, gp59, and gp60; Figure 4A). We performed a metagene analysis and found that across annotated PhiPA3 genes, ChmC shows enriched binding in a ∼50-bp region centered 25-30 bp downstream of the start codon (Figure 4C). We performed motif analysis to identify any defined sequences bound by ChmC, but could not unambiguously identify a preferred binding motif. Finally, we also observed high sequence coverage in short regions that do not correlate with annotated genes, including peaks between the *chmA* (gp53) and gp54 genes (Figure 4A) and between gp203 and gp204 (Figure S4A). These data suggest that PhiPA3 encodes small regulatory RNAs similarly to the related jumbo phage PhiKZ (30), and that ChmC binds many of these RNAs in addition to binding protein-coding mRNAs.

### ChmC knockdown halts jumbo phage infections and globally decreases phage protein levels

The jumbo phage nucleus prevents Cas9-based targeting of the phage genome due to its inaccessibility to host-encoded CRISPR/Cas enzymes (8, 9). Because phage mRNAs are transported into the host cytoplasm for translation, however, jumbo phages are susceptible to CRISPR-based translational knockdown strategies (8, 9). In related work, we used catalytically-dead *Ruminococcus flavefaciens* Cas13d (dCas13d) and a guide RNA overlapping the translation start site of an mRNA to efficiently inhibit translation of proteins encoded by the *E. coli* nucleus-forming phage Goslar; we term this method CRISPRi-ART (CRISPR interference by Antisense RNA Targeting) (31, 32) To determine the biological roles of ChmC, we designed guide RNAs that target the Goslar *chmC* gene (gp176) and found that when expressed alongside dCas13d, these guide RNAs block ChmC translation in infected cells as judged by western blotting (Figure S5A-B). Infection of *E. coli* MC1000 cells expressing dCas13d and *chmC*-targeting guide RNAs significantly reduced phage titer, with the most effective guide (guide 3) reducing the efficiency of plaquing to ∼4% of the efficiency observed in cells encoding a non-targeting guide RNA (Figure S5B-C). By microscopy, we observed that ChmC translational knockdown resulted in a strong reduction in phage bouquet formation in infected cells, indicative of a failure to assemble new virions (Figure S5D). Phage nuclei were also smaller and contained less DNA compared to control cells upon ChmC translational knockdown (Figure 5A, S5E-G). We could rescue these phenotypes by overexpressing a recoded ChmC resistant to dCas13d-mediated knockdown (Figure 5B, S6).

**Figure 5.**
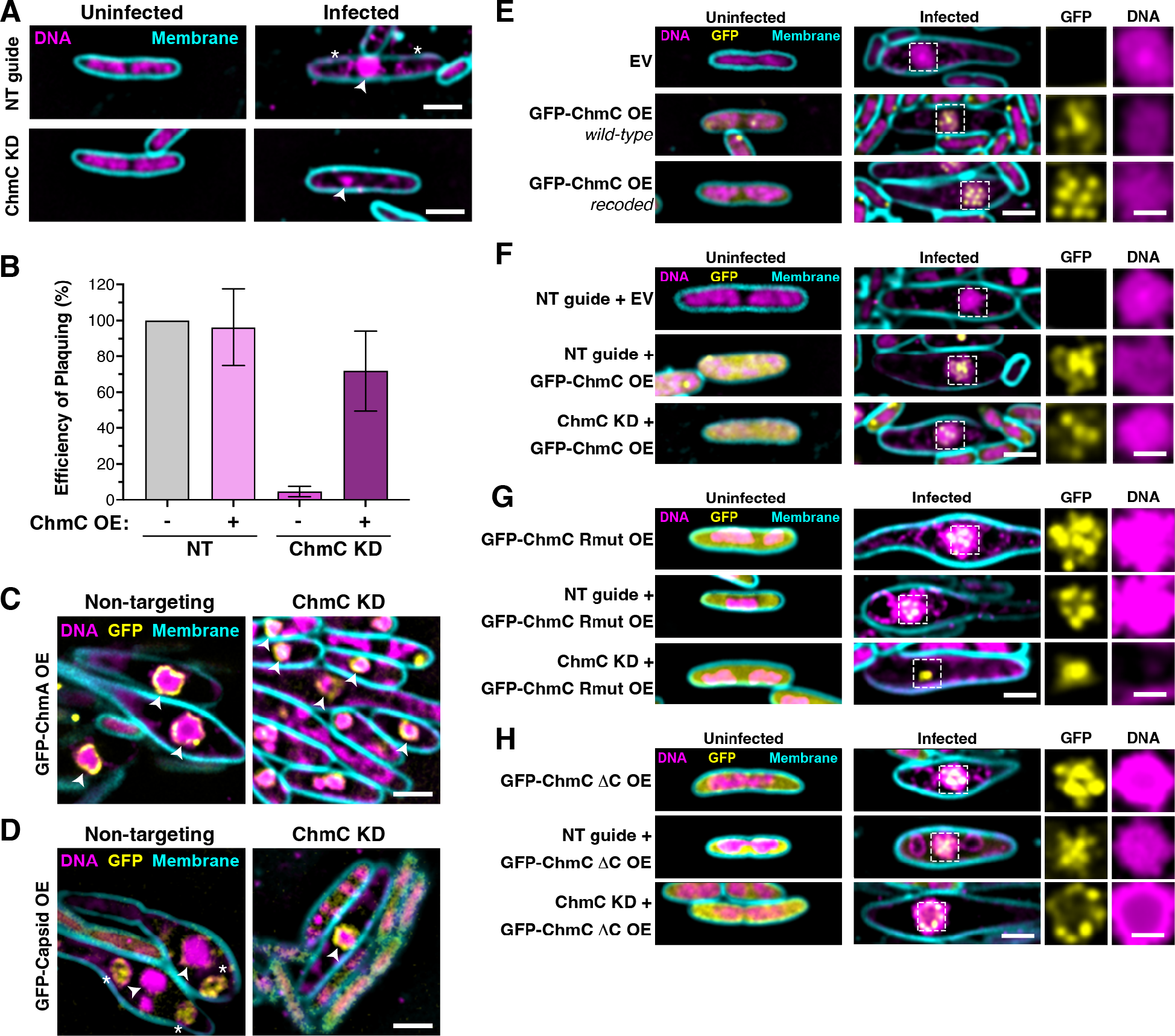
ChmC knockdown impairs phage nucleus development and infection progression. **(A)** Microscopy of *E. coli* MC1000 cells, either uninfected (left) or infected with Goslar (110 MPI; right). **(B)** Efficiency of plaquing of Goslar phage infecting *E. coli* MC1000 cells expressing dCas13d and either a non-targeting guide RNA (NT) or a guide RNA targeting the 5’ end of the ChmC (gp176) gene (guide 3; see **Figure S5** for analysis of three guide RNAs). Data are presented as mean +/-standard error of the mean of four biological replicates, calculated as a percentage of Goslar phage plaque forming units with no dCas13d/guide expression. See **Figure S6** for example plaque images and microscopy of knockdown/rescue cells. **(C)** Microscopy of *E. coli* MC1000 expressing GFP-tagged ChmA plus dCas13d and either a non-targeting guide RNA (left) or ChmC-targeting guide 3 (ChmC KD; right). DNA is stained with DAPI and shown in magenta; membranes are stained with FM4-64 and shown in cyan; GFP is shown in yellow. Arrowheads indicate the phage nucleus. Scale bar = 2 μm. See **Figure S7A** for further images. **(D)** Microscopy of *E. coli* MC1000 expressing GFP-tagged capsid protein plus dCas13d and either a non-targeting guide RNA (left) or ChmC-targeting guide 3 (ChmC KD; right). DNA is stained with DAPI and shown in magenta; membranes are stained with FM4-64 and shown in cyan; GFP is shown in yellow. Arrowheads indicate the phage nucleus, and asterisks indicate phage bouquets. See **Figure S7B** for further images. **(E)** Microscopy of *E. coli* MC1000 cells expressing no protein (EV, top row) or GFP-tagged ChmC (gp176, wild-type sequence in middle row and recoded sequence in bottom row), infected with Goslar (110 MPI). Yellow: GFP; magenta: DAPI nucleic acid; cyan: FM4-64 membrane dye. Scale bar = 2 μm for main panels, 1 μm for individual GFP and DNA channels representing zoomed views of the boxed regions. **(F)** Microscopy of *E. coli* MC1000 cells expressing dCas13d plus a non-targeting guide RNA (top and middle rows) or a ChmC-targeting guide (bottom roe), expressing either no protein (EV, top row) or GFP-tagged recoded ChmC (middle and bottom rows), and infected with Goslar (110 MPI). Yellow: GFP; magenta: DAPI nucleic acid; cyan: FM4-64 membrane dye. Scale bar = 2 μm for main panels, 1 μm for individual GFP and DNA channels representing zoomed views of the boxed regions. **(G)** As panel (E), except cells in the middle and bottom row are expressing ChmC Rmut (see **Figure S8** for Rmut design and analysis by Goslar plaquing assay). **(H)** As panel (E), except cells in the middle and bottom row are expressing ChmC ΔC (see **Figure S8** for ΔC design and analysis by Goslar plaquing assay).

We next imaged Goslar-infected *E. coli* cells expressing GFP-tagged ChmA (gp246) or the major capsid protein (gp41). Infected cells expressing dCas13d and a non-targeting guide RNA showed characteristic expansion of the cell diameter around the developing phage nucleus, which was labeled by GFP-tagged ChmA (Figure 5C, S7A). Infected cells expressing dCas13d and a *chmC*-targeting guide RNA showed little to no expansion of the cell diameter, and phage nuclei were markedly smaller than in control cells (Figure 5C, S7A). ChmC knockdown also caused infected cells to fail to form phage bouquets, and the major capsid protein was visibly attached to the nuclear shell, typical of early-stage infections when capsids localize to the nuclear shell for genomic DNA packaging (Figure 5D, S7B) (4, 11). Together, these data suggest that knockdown of ChmC significantly slows or halts Goslar infections at an early stage, preventing full maturation of the phage nucleus and production of viable phage progeny.

We next examined the localization of ectopically-expressed GFP-ChmC, both in the context of an unperturbed Goslar infection (Figure 5E) and with phage-encoded ChmC knocked down by dCas13d (Figure 5F). In both cases, we observed that GFP-ChmC localizes to the phage nucleus. In contrast to PhiPA3 ChmC, which localizes across the nuclear shell, Goslar ChmC forms discrete puncta both within the nucleus and along the nuclear shell. These puncta could represent hubs of transcription within the nucleus, hubs for translocation of phage mRNAs through the nuclear shell, or both. We generated mutants of Goslar ChmC analogous to the PhiPA3 ChmC-Rmut and ΔC mutants (Figure S8A-B). Both mutants failed to rescue Goslar phage titer in ChmC knockdowns, indicating that they cannot support the phage life cycle on their own (Figure S7C-E). In infected cells, both mutants localized as puncta within the phage nucleus in cells expressing a non-targeting guide RNA (Figure 5G-H). In cells expressing a *chmC*-targeting guide RNA, however, ChmC-Rmut localized as a single punctum that lacked any DNA (Figure 5G). We interpret this as indicative of a failure in nuclear shell assembly around the initially-injected phage genome, resulting in an aborted nuclear shell containing ChmC but with no DNA content. In contrast, ChmC-ΔC supported growth of the phage nucleus and localized as puncta on the perimeter of the phage nucleus (Figure 5H). Since ChmC-ΔC cannot form RNA-protein condensates but does retain RNA binding *in vitro*, its ability to form puncta at the nuclear shell suggests that these puncta represent hubs for mRNA translocation across the nuclear shell. Overall, these data support a model in which ChmC is important across *Chimalliviridae* to support proper assembly and maturation of the phage nuclear shell, and to support assembly of new virions.

### ChmC knockdown results in a global reduction of phage protein levels

Our data on ChmC mRNA binding and localization are consistent with roles in stabilizing phage mRNAs and/or promoting their translocation through the phage nuclear shell for translation. We tested the effects of ChmC knock-down on global phage protein levels 110 minutes post-infection using TMT-tagged mass spectrometry. Overall, we detected 180 of 247 annotated Goslar proteins across three samples (non-targeting guide RNA and two ChmC-targeting guide RNAs), 143 of which showed a reduction in protein levels after ChmC knockdown (Figure 6A, Table S4). Using a separate time-course mass spectrometry dataset of Goslar-infected cells (Figure 6B, Table S5), we divided the Goslar proteome into groups of proteins that first appear in early (30 minutes post infection), middle (60 minutes), or late (90 minutes) infections. Consistent with our related finding that early-expressed genes are predominantly transcribed by the virion RNA polymerase in a membrane-bound “early phage infection” (EPI) vesicle (7, 10, 31), we found that early proteins were largely unaffected by ChmC knockdown (Figure 6C). In contrast, middle- and late-expressed proteins were significantly suppressed upon ChmC knockdown (Figure 6C), consistent with the idea that these genes are transcribed by the non-virion RNA polymerase in the context of the phage nucleus.

**Figure 6.**
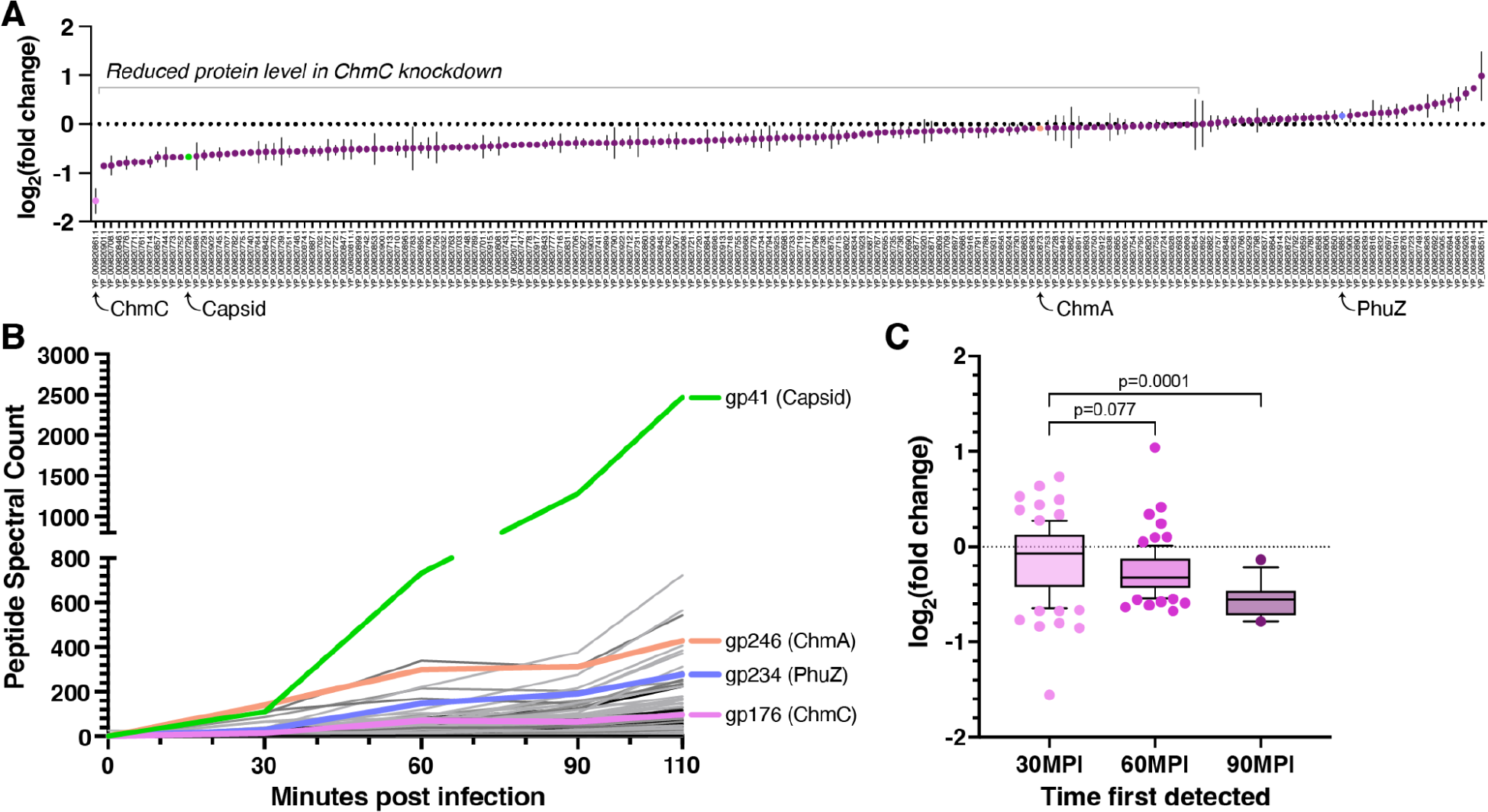
ChmC knockdown causes a global reduction in phage protein levels. **(A)** Log_2_(fold change) in expression of 180 Goslar proteins detected by TMT-tagged mass spectrometry analysis of Goslar-infected *E. coli* MC1000 cells (110 MPI) expressing dCas13d and *chmC*-targeting guide RNA 3 (see **Table S4** for full data, including for *chmC*-targeting guide RNA 2). Proteins are labeled by their NCBI accession numbers. ChmC (gp176), Capsid (gp41), ChmA (gp246), and PhuZ (gp234) are marked. **(B)** Spectral count of Goslar proteins from mass spectrometry of Goslar-infected *E. coli* MC1000 cell lysate at 0, 30, 60, 90, and 110 minutes post infection (MPI). ChmC (gp176) is shown in violet; Capsid (gp41) in green; ChmA (gp246) in salmon; and PhuZ (gp234) in blue. See **Table S5** for full data. **(C)** Aggregate log_2_(fold change) in expression upon ChmC knockdown (guide 3) for proteins first detected at 30 minutes (pink, n=84), 60 minutes (violet, n=74), and 90 minutes (purple, n=13) post Goslar infection. Box shows 25^th^ to 75^th^ percentile with mean value marked; whiskers show 10^th^ to 90^th^ percentile; and outlier points are shown individually. P values from one-way ANOVA comparisons are shown.

## Discussion

The nucleus-like compartment assembled by *Chimalliviridae* to protect their genomes from DNA-targeting host immune systems like restriction enzymes and CRISPR/Cas nucleases introduces major challenges to the phage life cycle, principally the need to translocate mRNAs out of the phage nucleus and to translocate specific phage proteins into the nucleus. Here, we identify an abundant and early-expressed protein, ChmC, that is conserved across *Chimalliviridae* and is encoded in a conserved block of genes alongside several subunits of the phage’s non-virion RNA polymerase. ChmC adopts a nucleic acid binding fold, binds phage mRNAs, and forms condensates with RNA in vitro. PhiPA3 ChmC localizes to the phage nuclear shell, while Goslar ChmC forms puncta that localize both within the phage nucleus and along the nuclear shell. Targeted knockdown of Goslar ChmC results in reduced phage nucleus size and a failure to form phage bouquets in infected cells, results in a global reduction of phage protein levels, and causes a dramatic reduction in viral plaque formation. Together, these data show that ChmC plays crucial roles in the life cycle of *Chimalliviridae*, potentially aiding mRNA transcription within the phage nucleus, promoting mRNA translocation through the nuclear shell, and/or directly promoting translation in the cytoplasm of infected cells.

Our structure predictions and biochemical characterization indicate that ChmC adopts a Whirly domain fold, first identified in single-stranded DNA/RNA binding proteins in plants where they are involved in transcriptional responses to stress (17, 33) . In bacteria, Whirly-related proteins are primarily involved in binding and compacting the nucleoid through their non-specific DNA binding activity (34–37). ChmC preferentially binds RNA over DNA *in vitro*, and shows enriched binding near the start codons of many phage mRNAs. While we do not detect any sequence motifs specifically recognized by ChmC, the protein may nonetheless specifically recognize particular mRNA sequences or structures that determine this binding pattern. Overall, these findings demonstrate that while ChmC is structurally related to other Whirly domain proteins, it has adopted distinct RNA regulatory roles in *Chimalliviridae*.

While bacteriophages generally do not encode RNA binding proteins, diverse eukaryotic viruses encode RNA binding proteins with diverse roles in the viral life cycle. Many such proteins also form RNA-protein condensates like ChmC (38). Coronaviruses like SARS, MERS, and SARS-CoV-2 encode an RNA-binding nucleocapsid (N) protein that promotes viral RNA production and suppresses host responses through its ability to form RNA-protein condensates, in addition to packaging viral genomic RNA into virions (39). Reminiscent of ChmC’s likely role in mRNA translocation through the nuclear shell, Influenza NEP (nuclear export protein) and HIV Rev are both RNA binding proteins that promote export of viral RNAs produced in the host-cell nucleus into the cytoplasm (40).

Our data show that ChmC is crucial for the proper progression of infection in *Chimalliviridae*, likely by aiding the translocation of phage mRNAs through the nuclear shell to promote translation. In *E. coli* cells infected with phage Goslar, ChmC forms puncta both within the phage nucleus and along the nuclear shell itself. These puncta are not simply RNA-protein condensates, since shell-associated puncta are also formed by ChmC-ΔC, which cannot form condensates with RNA. We hypothesize that these ChmC puncta may represent hubs for the translocation of mRNAs through the nuclear shell. These hubs are likely built around shell-penetrating pores, potentially the same ChmB pores that mediate capsid docking and genome packaging in late-stage infections (13). The location of *chmC* within a conserved block of genes encoding nvRNAP subunits further suggests a direct functional link between mRNA transcription and translocation through the nuclear shell, mediated by ChmC. Further work will be required to establish whether ChmB or other proteins are required for this process.

The phage nucleus of *Chimalliviridae* is a fascinating example of convergent evolution, representing a functional analog of the eukaryotic nucleus complete with physical segregation of the genome from the cytoplasm and specific mechanisms for mRNA export and protein import. Our work reveals the first identified RNA binding protein necessary for bacteriophage replication, with potential roles in mRNA production, export, and translation to support the unique life cycle of nucleus-forming jumbo phages.

## Acknowledgements

The authors thank Elizabeth Villa and members of the Corbett and Pogliano labs for helpful discussions. The authors acknowledge support from the National Institutes of Health (R01 GM129245 to J.P.; R35 GM144121 to K.D.C.; R01 HG004659 to G.W.Y.; S10 OD021724 for shared mass spectrometry resources). This research was supported in part by the Howard Hughes Medical Institute Emerging Pathogens Initiative (to J.P. and K.D.C.). J.A.D. is an Investigator of the Howard Hughes Medical Institute. Q.L. was supported by an individual predoctoral fellowship from the American Heart Association. B.A.A. was supported by m-CAFEs Microbial Community Analysis & Functional Evaluation in Soils, a Science Focus Area led by Lawrence Berkeley National Laboratory based upon work supported by the US Department of Energy, Office of Science, Office of Biological & Environmental Research [DE-AC02-05CH11231]. B.F.C. was supported by U. S. Department of Energy, Office of Science, through the Genomic Science Program, Office of Biological and Environmental Research, under the Secure Biosystems Design Initiative project Intrinsic Control for Genome and Transcriptome Editing in Communities (InCo-GenTEC). This paper was typeset with the bioRxiv word template by @Chrelli: www.github.com/chrelli/bioRxiv-word-template

## Materials and Methods

### Bacterial strains, growth conditions and phage preparations

For PhiPA3 phage, *P. aeruginosa* K2733 (PA01 efflux pump knockout; (*ΔMexAB-OprMΔMexCD-OprJΔMexEF-OprNΔMexXY-OprM*)) was used as the host. For Goslar phage, *E. coli* MC1000 (derived from *E*.*coli* MG1655) was used as the host. Both bacterial strains were cultured in Luria-Bertani (LB) media or LB top agar (0.35% agar) at 30°C (*P. aeruginosa*) or 37°C (*E. coli*). To amplify phages, 100 µL of liquid culture at OD_600_=0.6 was mixed with 20 µL of high-titer phage lysate and incubated at room temperature for 20 minutes, then the mixture was added to 5 mL of warm LB top agar, poured onto LB plates, and incubated overnight. The next day, 5 mL of Phage Buffer (10 mM Tris-HCl pH 7.5, 10 mM MgSO_4_, 68 mM NaCl, and 1 mM CaCl_2_) was added to each plate and incubated at room temperature for 5 hours. The phage buffers were then collected and lysates were centrifuged at 15,000 rpm for 10 minutes. Supernatants were stored at 4°C with 0.01% chloroform.

### Plasmid constructions and transformation

Genes of interest were PCR-amplified with 25 bp homology arms from high-titer phage lysates and ligated into respective plasmid backbones using NEBuilder HiFi DNA Assembly Cloning Kit (New England Biolabs). Recombinant plasmids were transformed into *E. coli* DH5α and plated on LB agar containing appropriate antibiotics (25 µg/mL gentamicin sulfate, 100 µg/mL ampicillin, 100 µg/mL spectinomycin, or 100 µg/mL chloramphenicol). After plasmids were confirmed by DNA sequencing, chemically competent organisms of interest were transformed and selected on LB plates with relevant antibiotics. Selected colonies were grown in LB media with the antibiotics and stored in 25% glycerol at -80°C.

### Fluorescence microscopy of single cell infections

1% agarose pads were prepared on concavity slides with desired inducing reagents of arabinose or IPTG. For imaging *P. aeruginosa*, pad mixes also contained FM4-64 (1 µg/mL) to stain cell membranes and DAPI (1 µg/mL) to stain DNA. Strains of interest were resuspended from overnight incubated LB plates into 25% LB to an OD_600_= 0.3. 5 µL of each resuspension was spotted on a concave slide and incubated in a humidor for 2 hours in 37°C (*E. coli* stains) or 30°C (*P. aeruginosa* strains). For phage infections, 10 µl of phages (10^10^ pfu/mL) were added to cells (resulting in a multiplicity of infection (MOI) of ∼7) and incubated until the desired time point. For imaging *E. coli*, dyes were added by spotting 7 µl of the mix containing (2 µg/mL DAPI, 4 µg/mL FM4-64, 25% LB). The slides were sealed with a coverslip and fluorescent microscopy was performed using a DeltaVision Spectris Deconvolution Microscope (Applied Precision). Regions of interests were imaged using at least 8 Z-axis stacks from the middle focal plane in 0.15 µm increments. Final images were created by DeltaVision SoftWoRx Image Analysis Program and its deconvolution algorithm and analyzed by Fiji ImageJ.

### Protein structure prediction

To model the structure of PhiPA3 or Goslar ChmC tetramers or PhiPA3 nvRNAP, we used AlphaFold multimer (14, 41) using ColabFold (42) installed locally on a Linux workstation with NVIDIA RTX 3090 GPU (https://github.com/YoshitakaMo/localcolabfold).

### Protein purification and characterization

For protein expression, *E. coli* strain Rosetta 2 (DE3) pLysS (EMD Millipore) cells were transformed with plasmids and grown overnight in LB plus appropriate antibiotics. The next day, cultures (1L 2XYT media plus antibiotics in 2L shaker flasks) were started and grown at 37°C until they reached an OD_600_ of 0.7, then induced with 0.25 mM IPTG and moved to 20°C for 16 hours. The cells were collected by centrifugation and resuspended in a buffer containing 25 mM Tris-HCl pH 7.5, 10% glycerol, 1 mM NaN_3_, 300 mM NaCl, 5 mM imidazole, and 5 mM β-mercaptoethanol. The proteins were purified using Ni^2+^ affinity chromatography (Ni-NTA agarose, Qiagen) and then passed over an anion-exchange column (Hitrap Q HP, Cytiva). Eluted fractions were concentrated and passed over a size exclusion column (Superdex 200, Cytiva) in GF buffer (Buffer A and 300 mM NaCl and 1 mM dithiothreitol). Fractions corresponding to the peak of interest were concentrated using ultrafiltration (Amicon Ultra, EMD Millipore) to reach a concentration of 10 mg/ml and stored at 4°C.

For analysis of molecular weight in solution using size exclusion chromatography coupled to multi-angle light scattering (SEC-MALS), 100 µl of purified protein at a concentration of 5 mg/ml was injected onto a size exclusion column (Superdex 200 Increase 10/300 GL, Cytiva) in GF buffer, then light scattering and refractive index profiles were collected using mini-DAWN TREOS and Optilab T-rEX detectors (Wyatt Technology). SEC-MALS data were analyzed using ASTRA software version 8.

### DNA and RNA binding assays

For measurement of DNA and RNA binding affinity by fluorescence polarization, 30 nM of a 22-base DNA (sequence ATTGTACCAC-TATTCCGAACAA) or RNA (sequence AUUGUACCAC-UAUUCCGAACAA) was mixed with the indicated concentration of purified PhiPA3 gp61 in FP buffer (20 mM HEPES pH 7.5, 75 mM KCl, 2 mM DTT, 5% glycerol, 0.02% NP40 substitute, 0.15 mg/mL BSA), and incubated for 10 minutes at room temperature. For RNA binding, reactions were supplemented with 1.25 mM RNaseOUT (ThermoFisher Scientific #10777019). Fluorescence polarization was measured with a Tecan Infinite M1000 PRO fluorescence reader, and data was analyzed by GraphPad Prism using a co-operative binding model.

### Knockdown of the phage proteins with dCas13d

31 nucleotide-long RNA guides that target the ribosome binding side or translational start of the gene of interest were designed and cloned into entry vectors with *Ruminococcus flavefaciens* Cas13d with mutations R239A, H244A, R858A, H863A (dCas13d). dCas13d is expressed under a tetR/tetA promoter, and guide RNAs are expressed under a J23119 promoter (31, 32). Plasmid selection was performed with chloramphenicol while appropriate anhydrotetracycline (aTc) concentration allows the expression of dCas13d. Plasmids were transformed to host MC1000 by electroporation and selected with 100 µg/mL chloramphenicol. For knockdown of ChmC in Goslar infections, *E. coli* strains were induced with 50 nM aTc and incubated at 37°C for 2 hours before infection.

### Western Blot

Samples were collected for western blot simultaneously with sample collection for TMT-tag mass spectrometry. Briefly, overnight liquid cultures of *E. coli* MC1000 strains were diluted to OD_600_=0.1 in fresh media with appropriate antibiotics and grown to OD_600_=0.6. 10 mL of pad mix (1% agarose, 25% LB, 50 nM aTc and 30 µg/mL chloramphenicol) was poured into a 6 cm petri dish. When the mixture solidified, 200 µL of each *E. coli* strain at OD_600_=0.1 was spread on a pad. Petri dishes were placed in a 37°C humidor and incubated for 2 hours. Cells were infected with 100 µL of Goslar phage (titer 10^9^ PFU/ml). At 0, 30, 60, and 90 minutes post infection cells were resuspended with 1 mL of 25% LB media and collected into 1.7 mL tubes, then pelleted by centrifugation at 4000 rpm at 4°C, then washed with 25% LB. Pellets were aliquoted into 5 tubes for the final wash, then washed pellets were stored at -80°C.

For cell lysis, samples were thawed on ice for 5 minutes and gently mixed with 500 µL of lysis buffer (10% Glycerol, 25 mM Tris pH 7.5, 150 mM NaCl supplemented with 4mg/mL lysozyme, 20µg/mL DNase I, 2x Complete Protease Inhibitor, 0.4 mM PMSF). Suspensions were incubated on ice for 1 hour, then sonicated for 30 seconds (Branson Sonifier) with duty cycle 40, output level 4 on ice. Suspensions were centrifuged at 15000 rpm for 30 minutes at 4°C. 40 µL of supernatant for each sample was mixed with 2x Laemmli buffer and boiled for 5 minutes at 95°C. 10 µL of each sample was loaded onto a 4–20% Mini-PROTEAN TGX Precast Protein Gel (Bio-Rad) and run at 180 V for 45 minutes. Proteins were transferred to PVDF membranes using a Trans-Blot Turbo RTA Mini 0.2 µm PVDF Transfer Kit (Bio-Rad) according to the manufacturer’s instructions, using a Trans-Blot Turbo Transfer System (Bio-Rad) at Turbo setting. Membranes were blocked (5% Non-Fat Dry Milk in TBST) for an hour in room temperature on a shaker. The solution was replaced with blocking buffer with appropriate dilutions of primary antibody (1:500 Rabbit anti-ChmC (Genscript, custom-generated) or 1:2,000 Rabbit anti-OmpA, (Joe Pogliano, custom-generated)) and incubated overnight on a shaker at 4°C. The next day, membranes were washed 3 times with TBS-T for 15 minutes and incubated with solution containing secondary antibody (1:10,000 HRP Goat anti-rabbit IgG, Thermo Fisher Scientific #65-6120) in blocking buffer at room temperature for 1 hour. The membrane was washed 3 times for 15 minutes each. For signal detection from HRP, Amersham ECL Select Western Blotting Detection Reagent (Cytiva) was used according to manufacturer’s instructions. Membranes were imaged with a ChemiDoc Imaging System (Bio-Rad) in Protein Blot - Chemiluminescence setting. Anti-OmpA membranes were imaged with 15 seconds exposure, while anti-ChmC membranes were imaged with 300 seconds exposure.

### Mass spectrometry of phage infections

For *P. aeruginosa*, overnight bacterial cultures in LB media were diluted to OD_600_=0.1 in fresh media, then further grown to OD_600_=0.5. Regrown cultures were diluted 1:10 into 50 mL total volume in 250 mL flasks and grown in LB supplemented with 0.2 mM CaCl_2_. Cells were infected with phage PhiPA3 at a multiplicity of infection (MOI) of 3 when they reached OD_600_=0.3, then collected at the indicated time points. Cultures were pelleted by centrifugation at 4000 rpm at 4°C. Cell pellets were washed with 25% LB to remove any free phage. After the last wash, pellets were snapfrozen in liquid nitrogen and stored at -80°C.

For *E. coli*, overnight cultures were diluted to OD_600_=0.1 in fresh media, then further grown to OD_600_=0.6. 10 mL of pad mix (1% agarose, 25% LB, 50 nM aTc and 30 ug/mL chloramphenicol) was poured into a 6 cm petri dish for each infection. When cultures reached desired OD_600_, each strain was prepared as 200 µL of OD_600_=0.1 and spread on the prepared pads. Petri dishes were placed in a 37°C humidor and incubated for 2 hours, then 100 µL of Goslar (titer 10^9^ PFU/ml) was spread on the pad and incubated at 37°C for the indicated times. At the time of collection, 1 mL of 25% LB media was added and cells were carefully resuspended. Cells were pelleted by centrifugation at 4000 rpm at 4°C, washed with 25% LB, then snapfrozen and stored at -80°C for mass spectrometry.

### Plaque Assays for Phage Infectivity

For plaque assays, 500 µl of saturated overnight bacterial cultures were mixed with 4.5 ml of 0.35% LB top agar. This mixture was poured on an LB plate that contains 100 µg/mL chloramphenicol and 50 nM aTc. After 30 minutes of incubation at room temperature for the mixture to dry, 3 µl of 10-fold serial dilutions of Goslar lysates (10^9^ pfu/mL) were spotted on each plate. Plates were incubated at 37°C for 16 hours, then plaques were counted.

### GFP pulldowns

GFP pulldowns were performed with GFP-Trap Magnetic Agarose beads (Proteintech). *P. aeruginosa* strains carrying pHERD30T plasmids expressing GFP-tagged proteins of interest were grown in LB with 25 µg/mL gentamicin sulfate. Saturated overnight cultures were diluted to OD_600_=0.1 in fresh media, then grown at 30°C until they reached OD_600_=0.5. Cultures were diluted 1:10 in 50 mL LB supplemented with arabinose, gentamicin sulfate, and calcium chloride. Once the cells reached at OD_600_=0.3, they were infected with PhiPA3 at MOI 3. The cultures were collected at 45 minutes post-infection, centrifuged, and the resulting cell pellets were stored at -80°C.

To perform the GFP pulldown, frozen cell pellets were thawed and incubated for 1 hour with 500 µL lysis buffer (10% glycerol, 25 mM Tris-HCl pH 7.5, 150 mM NaCl, 4 mg/mL lysozyme, 20 µg/mL DNase I, 2x cOmplete Protease Inhibitor, 0.4 mM PMSF). The cell suspensions were then sonicated (10 rounds x 20 pulses/round, Duty Cycle 40, Output 4), and the resulting lysed cells were centrifuged (30 minutes at 15,000 rpm at 4ºC). Beads were washed 3 times with a wash buffer (10 mM Tris-HCl pH 7.5, 150 mM NaCl, 0.5 mM EDTA) and added to the supernatant of the cell lysate. The mixture was rotated end-to-end for 1 hour at 4°C. The beads incubated in the cell lysate were washed 3 times with 1 mL wash buffer. After the last wash, beads were stored at -80°C for later analysis by SDS-PAGE and mass spectrometry.

### eCLIP-Seq

Overnight bacterial cultures in LB media were diluted to OD_600_=0.1 and grown to OD_600_=0.5 at 30°C. The cultures were diluted 1:10 into 50 mL total volume in 250 mL flasks and grown in LB supplemented with 0.2 mM CaCl_2_ and 0.1% Arabinose. Cells were infected with phage PhiPA3 at a multiplicity of infection (MOI) of 3 when they reached OD_600_=0.3. After 45 minutes of infection, cultures were collected and centrifuged with 4000 rpm at 4°C for 8 minutes. Cells were washed with PBS 2 times and resuspended in 10mL fresh PBS. The samples were spread on 10 cm petri-dish to coat the surface and UV crosslinking was performed with 400 mJ/cm2 at 254 nm.

Samples were collected from the petri dish and pelleted at 4C at 4000 rpm for 10 minutes and snap-frozen in liquid nitrogen to be stored at -80°C.

Each sample was treated with immunoprecipitation protocol with GFP-Trap beads. After the RNA-bound proteins were collected, cells were treated with FastAP (ThermoFisher) and T4 PNK (NEB), then barcoded RNA adapters were ligated to the 3′ end (T4 RNA Ligase, NEB). Samples were separated by SDS-PAGE and transferred to nitrocellulose membranes. The regions corresponding to the approximately expected size of sfGFP-alone and gp61-sfGFP were excised, and the membrane was suspended in a buffer with proteinase K (NEB). RNA isolation was performed with phenol/chloroform extraction and purified on spin columns (Zymo Research). Reverse-transcription was performed with AffinityScript (Agilent). cDNAs were treated with ExoSAP-IT (Affymetrix) to remove the excess oligonucleotides. Second DNA adapters (containing 5 [N5] or 10 [N10] random bases at the 5′-end) were ligated to the 5′-end of the cDNA (T4 RNA Ligase, NEB). The DNA was amplified by PCR and purified with PippinPrep system (Sage Science) and sequenced with Illumina HiSeq 4000. Libraries were analyzed for fragment size distribution on a D1000 Screentape (Agilent). Reads were processed and mapped to the PhiPA3 genome. Normalization of the eCLIP data was performed using the input samples prior to pulldown.

Reads were processed according to the protocol as previously described (43). Briefly, sequenced reads from both IP and corresponding size-matched input (SMInput) were trimmed of adapters and mapped to repeat elements first to remove non-uniquely mapped reads, then to the genome composed of both *P. aeruginosa* PA01 (NCBI RefSeq #GCF_000006765.1) and PhiPA3 (NCBI RefSeq #GCF_001502095.1). PCR collapsing was then performed, and CLIPper (44) was used to call peak clusters on each set of these uniquely mapped, deduplicated (usable) reads. Annotations from *P. aeruginosa* PA01 and PhiPA3 were used to construct a custom CLIPper index. Reads within these clusters were normalized against the SMinput sample using scripts (available at https://github.com/yeolab/eclip), and peaks found to be enriched above a log_2_(fold change) of 3 and -log_10_(Fisher Exact or Chi-square p-value) threshold of 3 were deemed significant and merged using IDR (45) to produce a set of reproducible peaks from replicates. Metagene plots were generated from usable reads, using a strategy laid forth by (46). Briefly, read densities from SMInput samples were subtracted from corresponding IP signals across the set of annotated start codons of expressed phage genes, using SMInput data as a proxy for gene expression.

### Protein identification by Mass Spectrometry

Frozen cell pellets were thawed and resuspended in 100 µL water. 10 µL of resuspended cells were mixed with 200 µL of 6M guanidine-HCl, vortexed and subjected to 3 cycles of 100°C for 5 minutes followed by cooling to room temperature. Boiled cell lysates were mixed with 1.8 mL of pure methanol and incubated at -20°C for 20 minutes. The mixture was centrifuged at 14000 rpm for 10 minutes at 4°C. All liquid was removed and pellet was resuspended in 200 µL of 8 M urea in 0.2 M ammonium bicarbonate and incubated at 37°C for 1 hour with constant agitation. 4 µL of 500 mM TCEP (Tris(2-carboxyethyl) phosphine) and 20 µL 400 mM chloro-acetamide were added to the samples.

Protein concentration was measured by BCA assay and 600 µL of 200 mM ammonium bicarbonate was added to bring the urea concentration to 2 M. 1 µg of sequencing-grade trypsin was added for each 100 µg of protein in the sample and incubated at 42°C for overnight. The next day, 50 µL of 50% formic acid was added (final pH=2), then samples were desalted with C18 solid phase extraction (Waters Sep-Pak C18 12 cc Vac Cartridge # WAT036915) according to the manufacturer’s protocol. Samples were resuspended in 1 ml phosphate-buffered saline and peptide concentration of each sample was measured with BCA (bicinchoninic acid assay).

Trypsin-digested peptides were analyzed by ultra-high pressure liquid chromatography (UPLC) coupled with tandem mass spectroscopy (LC-MS/MS) using nano-spray ionization. Nanospray ionization was performed with Orbitrap fusion Lumos hybrid mass spectrometer (Thermo) interfaced with nano-scale reverse-phase UPLC (Thermo Dionex UltiMate 3000 RSLC nano System) using a 25 cm, 75-micron ID glass capillary packed with 1.7-µm C18 (130) BEH beads (Waters corporation). Peptides transferred from C18 column into the mass spectrometer by a linear gradient (5–80% buffer B) using buffers A (98% H2O, 2% acetonitrile, 0.1% formic acid) and B (100% acetonitrile, 0.1% formic acid) at a flow rate of 375 µl/min over 3 hours. Mass spectrometer parameters were; MS1 survey scan using the orbitrap detector (mass range (m/z): 400-1500 (using quadrupole isolation), 120000 resolution setting, spray voltage of 2200 V, Ion transfer tube temperature of 275°C, AGC target of 400000, and maximum injection time of 50 ms) which was followed by a data dependent scans (top speed for most intense ions, with charge state set to only include +2-5 ions, and 5 second exclusion time, while selecting ions with minimal intensities of 50000 at in which the collision event was carried out in the high energy collision cell (HCD Collision Energy of 30%), and the fragment masses were analyzed in the ion trap mass analyzer (With ion trap scan rate of turbo, first mass m/z was 100, AGC Target 5000 and maximum injection time of 35 ms). Protein identification quantifications were carried out using Peaks Studio X (Bioinformatics Solutions Inc).

### TMT-tag Mass Spectrometry

Cell pellets from 200 µL of culture were resuspended in 600 µl of 8 M Urea in 100 mM Tris-HCl pH 8.0 and vortexed for 10 minutes. TCEP (tris(2-carboxyethyl)phosphine) was added to a final concentration of 10 mM, and samples were incubated at -20°C overnight to solubilize proteins. The next day, samples were vortexed until the solution was clear. Chloro-acetamide was added to the final concentration of 40 mM, and the mixture was vortexed for 5 minutes. Urea concentration was reduced to 4 M by adding an equal volume of 50 mM Tris-HCl pH 8.0 to the samples, then protein concentration was measured. LysC was added at 1:500 (w:w) LysC:protein ratio, and mixtures were incubated at 37°C on a roller for 6 hours. Urea concentration was further reduced to 2 M by adding 50 mM Tris-HCl pH 8.0. A 1:50 (w:w) ratio of trypsin was next added, and samples were incubated overnight. The next day, samples were acidified by adding TFA (trifluoroacetic acid) to 0.5% final concentration, and mixtures were vortexed for 5 minutes. Samples were centrifuged at 15,000 x g for 5 minutes to obtain aqueous and organic phases. The lower aqueous phase was collected and desalted with 100 mg C18-StageTips (Thermo Scientific) according to the manufacturer protocol. Samples were resuspended in Thermo Fisher iTRAQ dissolution buffer and peptide concentrations were measured by BCA. For TMT labeling, TMTpro 16plex (Thermo Scientific, A44520) was used according to the manufacturer protocol. For high pH fractionation, High pH Reversed-Phase Peptide Fractionation Kit (Pierce #84868) was used according to the manufacturer protocol.

Each fraction was analyzed by ultra-high-pressure liquid chromatography (UPLC) coupled with tandem mass spectroscopy (LC-MS/MS) and nano-spray ionization with an Orbitrap fusion Lumos hybrid mass spectrometer (Thermo Fisher Scientific) interfaced with nano-scale reversed-phase UPLC (Thermo Fisher Scientific Dionex UltiMate 3000 RSLC nano System). In these experiments, a 25 cm, 75-micron ID glass capillary packed with 1.7-µm C18 (130) BEH beads (Waters) were used. From C18 into the column, a linear gradient (5–80%) of acetonitrile at a flow rate of 375 µl/min was used to elute peptides into the mass spectrometer for 180 minutes. The acetonitrile gradient was created from Buffer A (98% H_2_O, 2% acetonitrile, 0.1% formic acid) to Buffer B (100% acetonitrile, 0.1% formic acid). Parameters of MS1 survey scan using the orbitrap detector (mass range (m/z): 400-1500 (using quadrupole isolation), 60000 resolution setting, spray voltage of 2200 V, Ion transfer tube temperature of 275 C, AGC target of 400000, and maximum injection time of 50 ms) were used for mass spectrometer. This was followed by data dependent scans (top speed for most intense ions, with charge state set to only include +2-5 ions, and 5 second exclusion time). Ions with minimal 50000 intensity were selected with a high energy collision cell (HCD Collision Energy of 38%) and the first quadrupole isolation window was set at 0.7 (m/z). Orbi-trap mass analyzers were used for analyzing the fragment masses (Ion trap scan rate of turbo, first mass m/z was 100, AGC Target 20000 and maximum injection time of 22ms). Peaks Studio X (Bioinformatic Solutions Inc.) was used for protein identification and quantification.

### Condensate analysis

Macromolecular condensation assays were conducted in vitro using phase separation buffer (20 mM HEPES pH 7.4, 50 mM NaCl) at a temperature of 25°C, a protein concentration of 30 µM, and an RNA concentration of 2 µM (for samples including RNA). To prepare samples, unlabeled gp61 was pre-mixed with Cy5-labeled gp61 (linked to an engineered N-terminal cysteine using maleimide linkage) at a ratio of 1:10 and diluted to 60 µM in phase separation buffer. For samples without RNA co-incubation, an equal volume of phase separation buffer was added to each protein sample to reach the final working concentration of 30 µM. For samples with RNA coincubation, an equal volume of RNA at 2X final concentration (4 µM for 40 base RNA; 166 nM for 2.3 kb RNA) in phase separation buffer was added and mixed gently. Samples were mixed in protein LoBind tubes (Eppendorf) and then immediately transferred into a 96-well non-binding plate (Greiner Bio-one). Samples were imaged immediately after transfer into the 96-well plate using a CQ1 confocal quantitative image microscope (Yokogawa) with a 20x-PH objective. The fluorescent signal was captured under a laser at 640 nm. For live imaging, the entire field was automatically captured every 10 minutes. For quantitation, condensates were identified by particle analysis in ImageJ (47). After thresholding, individual particles were counted and their areas measured. For each sample showing particles, the coefficient of variation was calculated as the standard deviation of particle area divided by the mean particle area.

**Figure S1.**
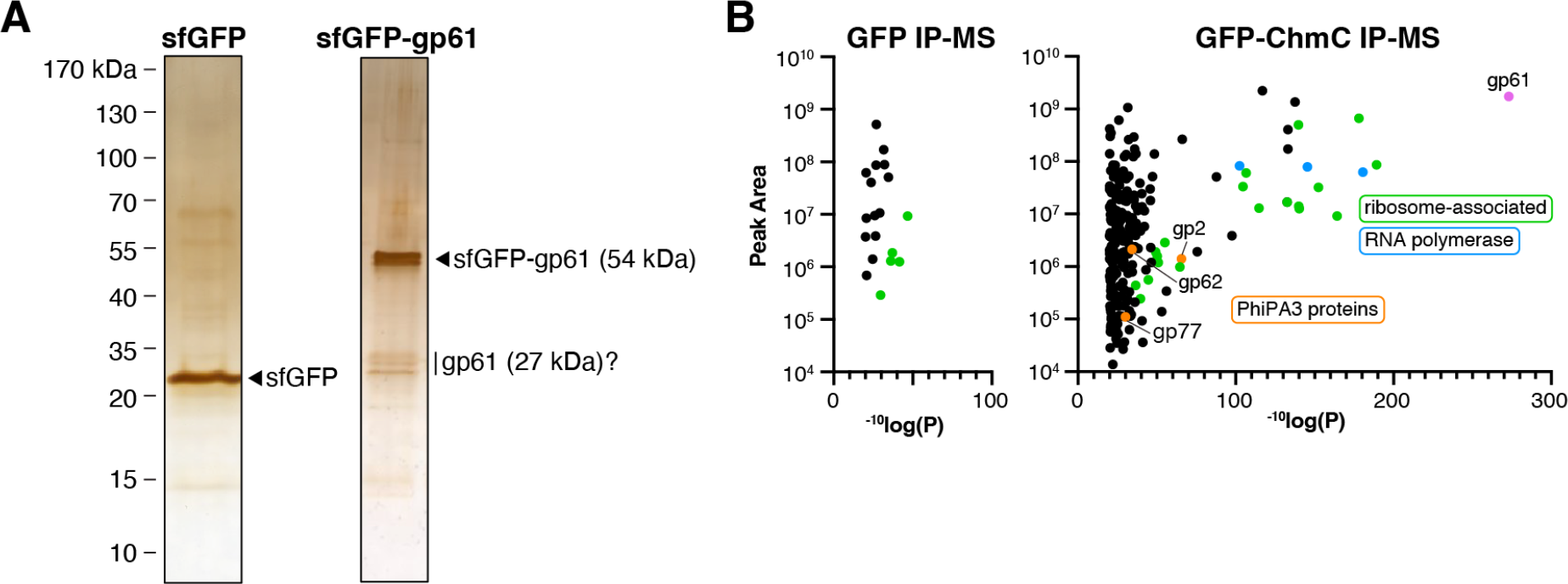
IP-MS of PhiPA3 gp61. **(A)** Silver-stained SDS-PAGE analysis of sfGFP (left) and sfGFP-gp61 (right) pulldowns from PhiPA3-infected *P. aeruginosa* cells. Tentative assignments of prominent bands are shown for each. **(B)** Mass spectrometry proteomics analysis (total peak area for assigned peptides per protein vs. ^-10^log(P) for sfGFP (left panel) versus sfGFP-fused gp61 (right panel) immunopurified from PhiPA3-infected *P. aeruginosa* (45 minutes post infection). gp61 is labeled and shown in pink, other PhiPA3 proteins are shown in orange and individually labeled, host ribosomal or ribosome-associated proteins are shown in green, and host RNA polymerase subunits are shown in blue. See **Table S3** for full mass spectrometry data.

**Figure S2.**
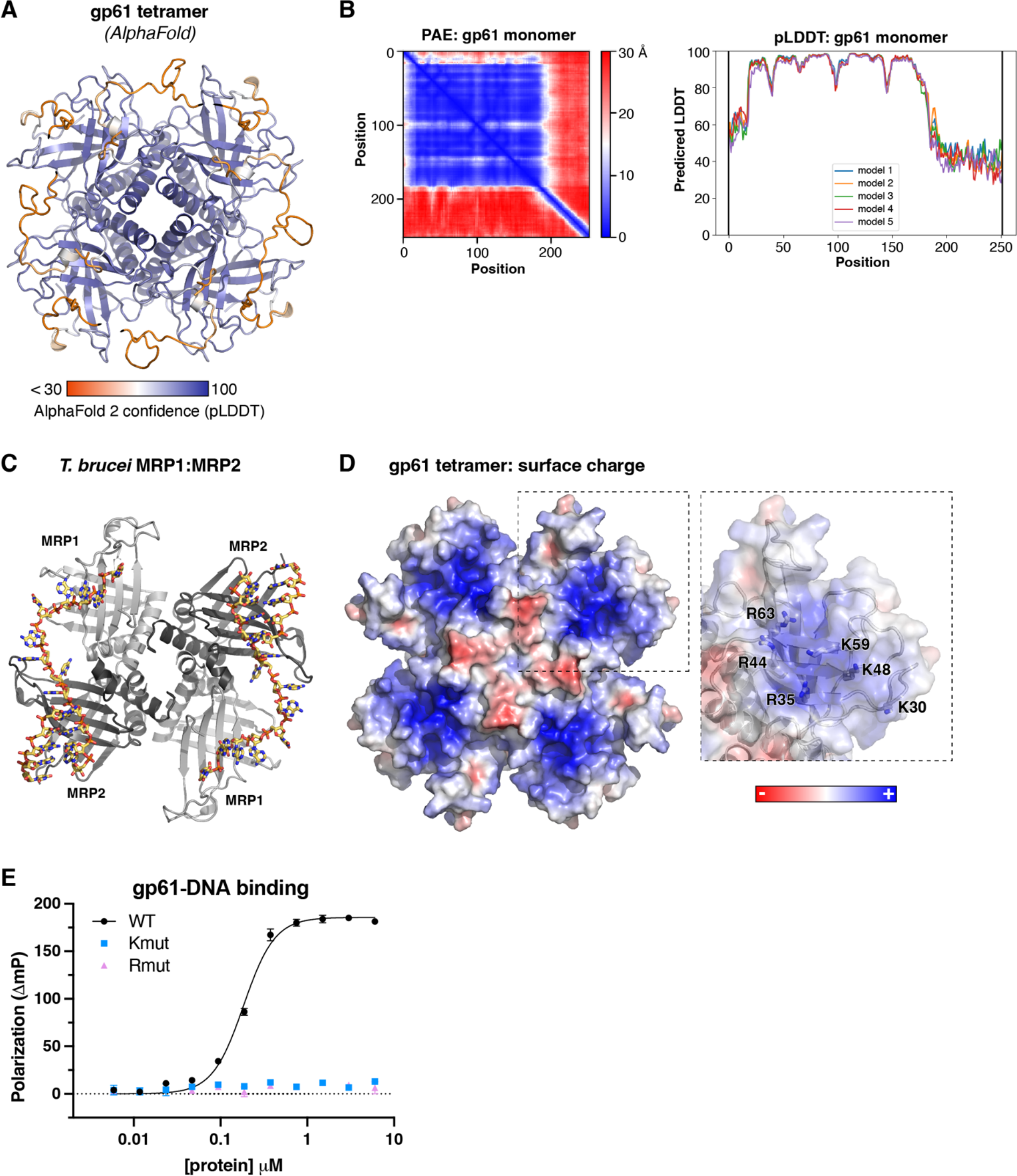
ChmC adopts an RNA-binding Whirly domain fold. **(A)** AlphaFold 2 predicted structure of a homotetramer of PhiPA3 ChmC (gp61), colored by confidence (pLDDT). **(B)** Predicted Aligned Error (PAE) and predicted local distance difference test (pLDDT) plots for one PhiPA3 ChmC monomer, as output by AlphaFold 2. **(C)** Structure of *T. brucei* MRP1:MRP2 bound to RNA (PDB ID 2GJE) (16). **(D)** Surface charge distribution of the AlphaFold 2 predicted structure of a homotetramer of PhiPA3 ChmC, calculated by APBS (46) in PyMOL version 2. **(E)** Single-stranded RNA binding of PhiPA3 ChmC wild-type (black circles, reproduced from **Figure 2D**), Kmut (blue squares), and Rmut (pink triangles), as measured by fluorescence polarization. Data points are shown as average +/-standard deviation of triplicate technical replicates.

**Figure S3.**
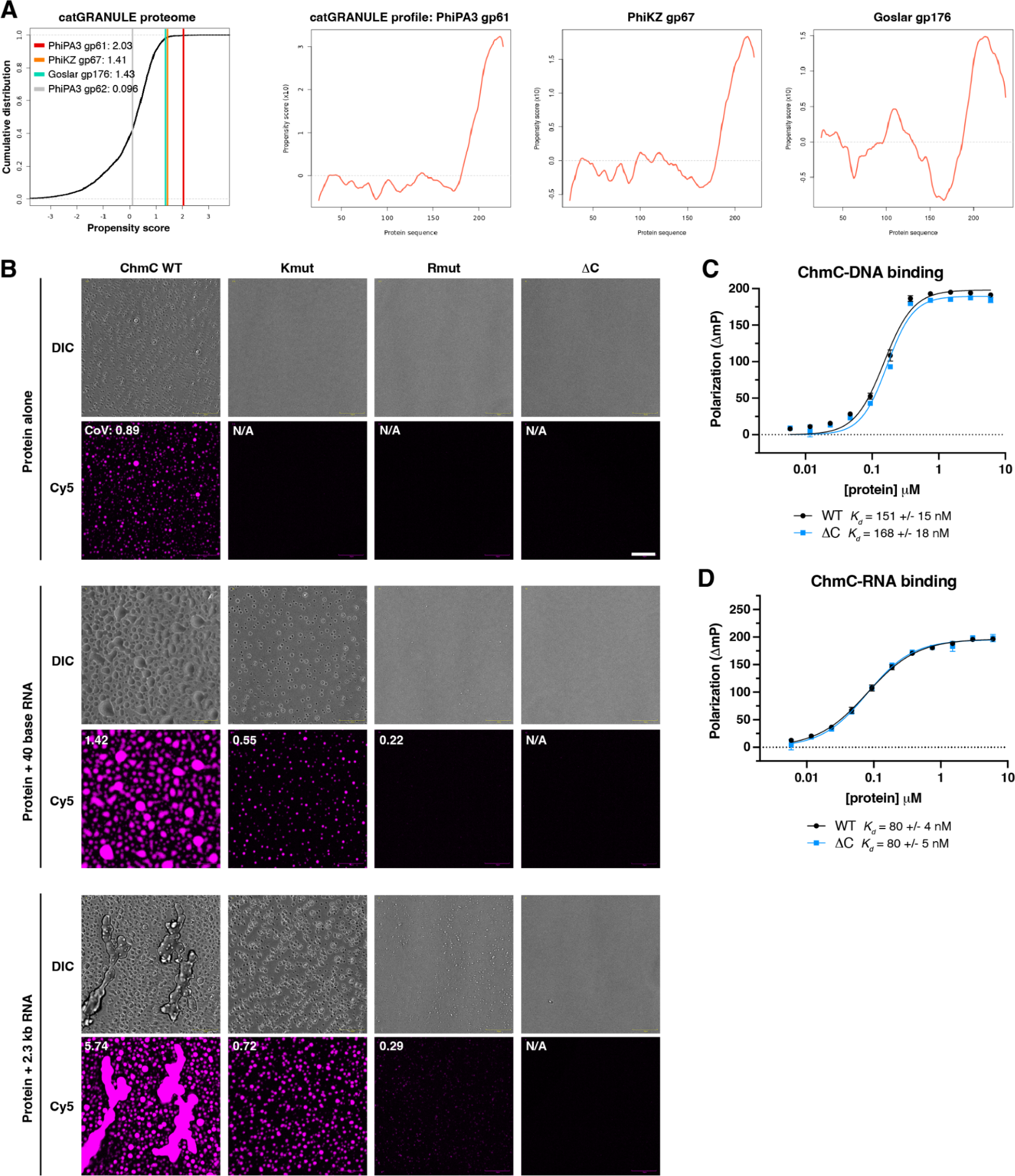
ChmC forms phase-separated condensates with RNA. **(A)** *Left:* catGRANULE output for ChmC from PhiPA3 (gp61), PhiKZ (gp67; 56% identical to PhiPA3 gp61), and Goslar (gp176; 30% identical to PhiPA3 gp61), plus a soluble protein (PhiPA3 gp62, a nvRNAP subunit) as a negative control. *Right:* catGRANULE profiles for ChmC from PhiPA3 (gp61), PhiKZ (gp67), and Goslar (gp176), showing that the disordered C-terminus of each protein likely drives phase separation. In agreement with catGRANULE output, the machine learning based PSPredictor server (47) strongly predicted condensation behavior for all three proteins (scores of 0.8706 for PhiPA3 gp61; 0.6531 for PhiKZ gp67; and 0.8461 for Goslar gp176 on a scale of 0 (no condensation predicted) to 1 (highly confident condensation behavior predicted)). These predictions were completely dependent on the C-terminal disordered regions, with removal of these regions resulting in low scores for all three proteins (0.0048 for PhiPA3 gp61 residues 1-188; 0.0024 for PhiKZ gp67 residues 1-187; and 0.0226 for Goslar gp176 residues 1-193). **(B)** Differential interference contrast (DIC) and fluorescence microscopy (Cy5) imaging of PhiPA3 ChmC (wild type, Kmut, Rmut, or ΔC; 10% Cy5-labeled) at 30 μM protein concentration, either alone (top row) or with 2 μM of a 40-base RNA (2.8 μg/mL; middle row) or 83 nM of a 2.3 kb RNA (5.8 μg/mL; bottom row). All images were taken 30 minutes after final dilution and mixing with RNA. For all conditions that showed condensate formation, the coefficient of variation (CoV) was calculated as the standard deviation of particle area divided by the mean particle area (WT protein alone n=423; WT + 40 base RNA n=222; Kmut + 40 base RNA n=247; Rmut + 40 base RNA n=5; WT + 2.3 kb RNA n=552; Kmut + 2.3 kb RNA n=368; Rmut + 2.3 kb RNA n=147). Scale bar = 30 μm. See Figure S2E for DIC imaging. **(C)** Single-stranded DNA binding of PhiPA3 ChmC WT (black squares) and ΔC (blue circles) as measured by fluorescence polarization. Data points are shown as average +/-standard deviation of triplicate technical replicates, and curves are fit with a cooperative binding model (Hill coefficient for WT ChmC binding DNA 2.0 +/-0.3; Hill coefficient for ChmC ΔC binding DNA 2.2 +/-0.4). **(D)** Single-stranded RNA binding of PhiPA3 ChmC WT (black squares) and ΔC (blue circles) as measured by fluorescence polarization. Data points are shown as average +/-standard deviation of triplicate technical replicates, and curves are fit with a cooperative binding model (Hill coefficient for WT ChmC binding RNA 1.2 +/-0.1; Hill coefficient for ChmC ΔC binding RNA 1.3 +/-0.1).

**Figure S4.**
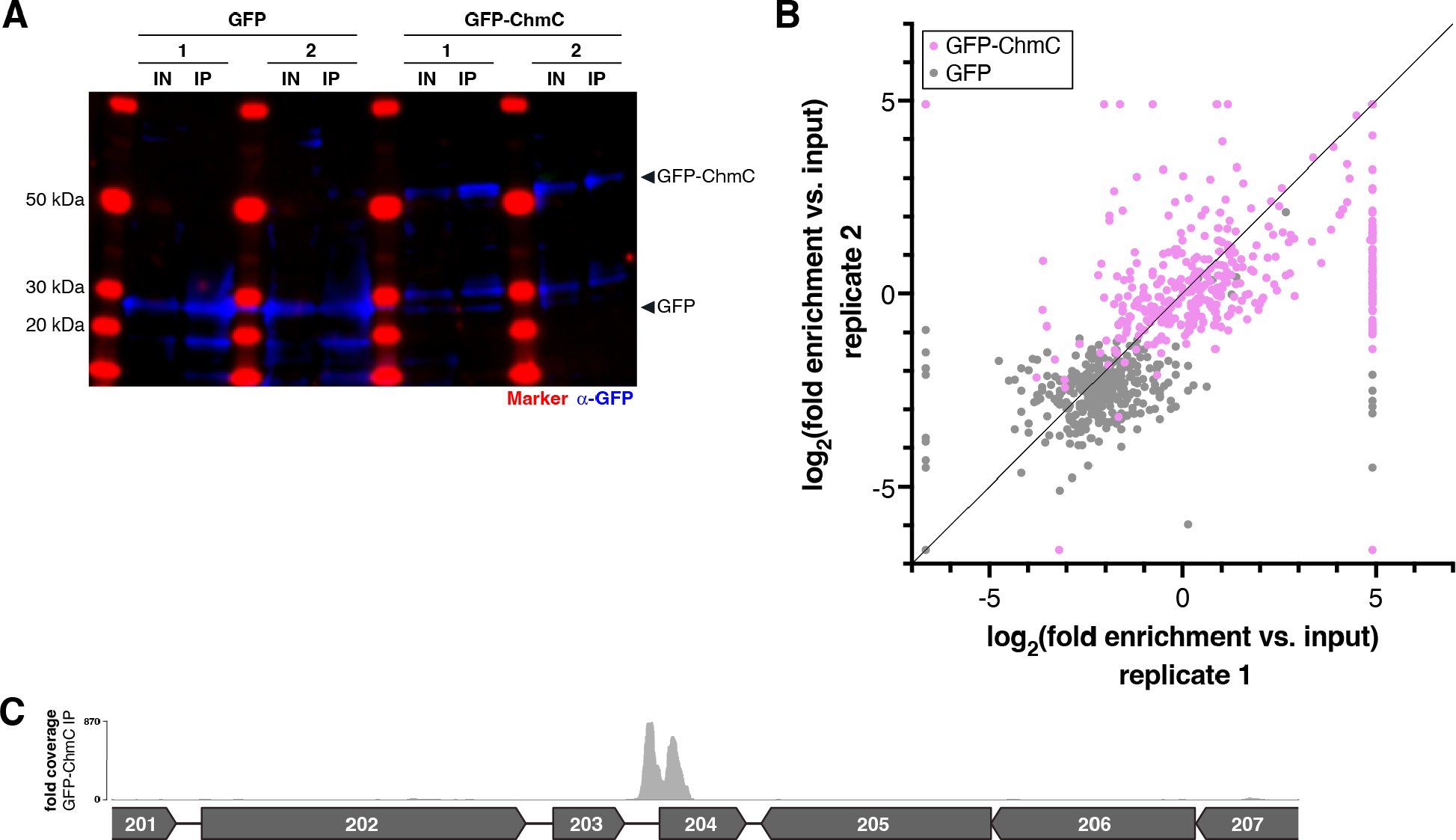
eCLIP analysis of ChmC-RNA interactions. **(A)** Western blot showing levels of GFP (blue) in eCLIP input (IN) and immunopurified (IP) samples. Replicates 1 and 2 for each sample (GFP or GFP-ChmC) are labeled. Molecular weight markers are shown in red and labeled. **(B)** Reproducibility of two replicates for eCLIP analysis of GFP (gray dots) and GFP-ChmC (pink dots), shown as log_2_(fold enrichment vs. input) for each phage gene. **(C)** Fold sequence coverage from eCLIP analysis of GFP-ChmC, showing large peaks in the region upstream of PhiPA3 gp204 and downstream of the gp204 start codon.

**Figure S5.**
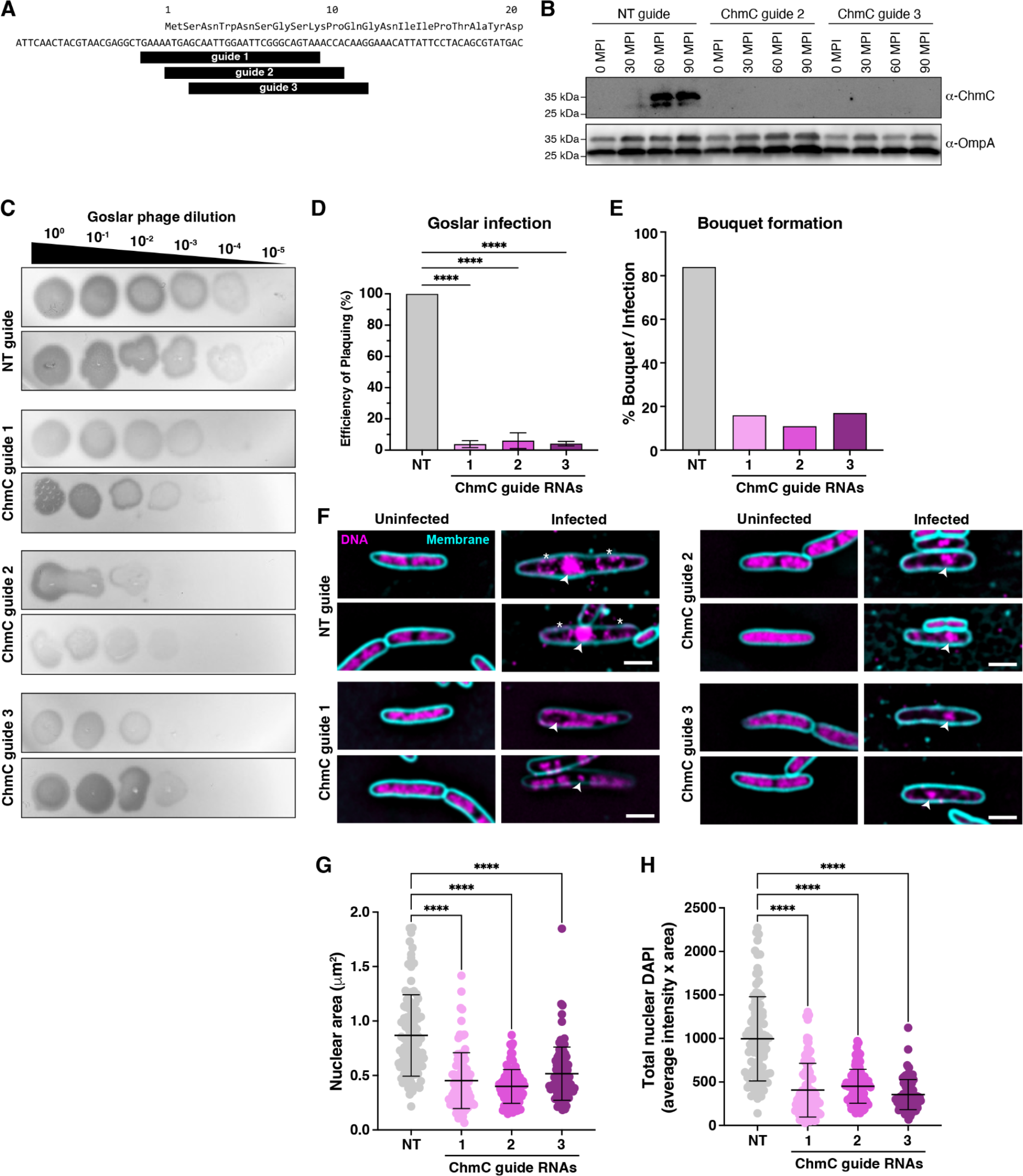
ChmC knockdown disrupts Goslar phage infections. **(A)** Design of three guide RNAs targeting the start codon of phage Goslar ChmC (gp176). **(B)** Western blot showing expression of ChmC in *E. coli* MC1000 cells expressing dCas13d and either a non-targeting guide RNA (NT) or guide RNAs targeting the 5’ end of the ChmC (gp176) gene (guide 2 or guide 3), upon high-multiplicity infection with phage Goslar. MPI: minutes post infection. α-ChmC indicates western blot with a custom-generated anti-ChmC antibody; α-OmpA indicates western blot with anti-*E. coli* OmpA antibody (loading control). **(C)** Plaque assays with phage Goslar on *E. coli* MC1000 cells expressing dCas13d and either a non-targeting guide RNA (NT) or one of three guide RNAs targeting the 5’ end of the ChmC (gp176) gene (guide 1, guide 2, or guide 3). **(D)** Efficiency of plaquing of phage Goslar on *E. coli* MC1000 cells dCas13d and either a non-targeting guide RNA (NT) or one of three guide RNAs targeting the 5’ end of the ChmC (gp176) gene (guide 1, guide 2, or guide 3). *** indicates a P value < 0.0002 in a one-way ANOVA test of statistical significance. **(E)** Percent of Goslar-infected *E. coli* MC1000 cells forming phage bouquets at 110 MPI. **(F)** Microscopy of *E. coli* MC1000 (uninfected at left; 110 MPI Goslar-infected at right) expressing dCas13d and either a non-targeting guide RNA (top) or one of three guide RNAs targeting the 5’ end of the ChmC (gp176) gene (guide 1, guide 2, or guide 3). DNA is stained with DAPI and shown in magenta; membranes are stained with FM4-64 and shown in cyan. Arrowheads indicate the phage nucleus, and asterisks indicate phage bouquets. Scale bar = 2 μm. **(G)** Phage nucleus area from 110 MPI Goslar-infected *E. coli* MC1000 cells expressing dCas13d and either a non-targeting guide RNA (NT; gray) or a guide RNA targeting the 5’ end of the ChmC (gp176) gene (guide 1 pink, guide 2 violet, guide 3 purple). Individual data points are shown, and error bars indicate mean +/-standard deviation. **** indicates a P value of <0.0001 in a one-way ANOVA test of statistical significance. **(H)** Total DAPI signal within the phage nucleus (calculated as the phage nuclear area multiplied by the average DAPI intensity in the phage nucleus) from 110 MPI Goslar-infected *E. coli* MC1000 cells expressing dCas13d and either a non-targeting guide RNA (NT; gray) or a guide RNA targeting the 5’ end of the ChmC (gp176) gene (guide 1 pink, guide 2 violet, guide 3 purple). Individual data points are shown, and error bars indicate mean +/-standard deviation. **** indicates a P value of <0.0001 in a one-way ANOVA test of statistical significance.

**Figure S6.**
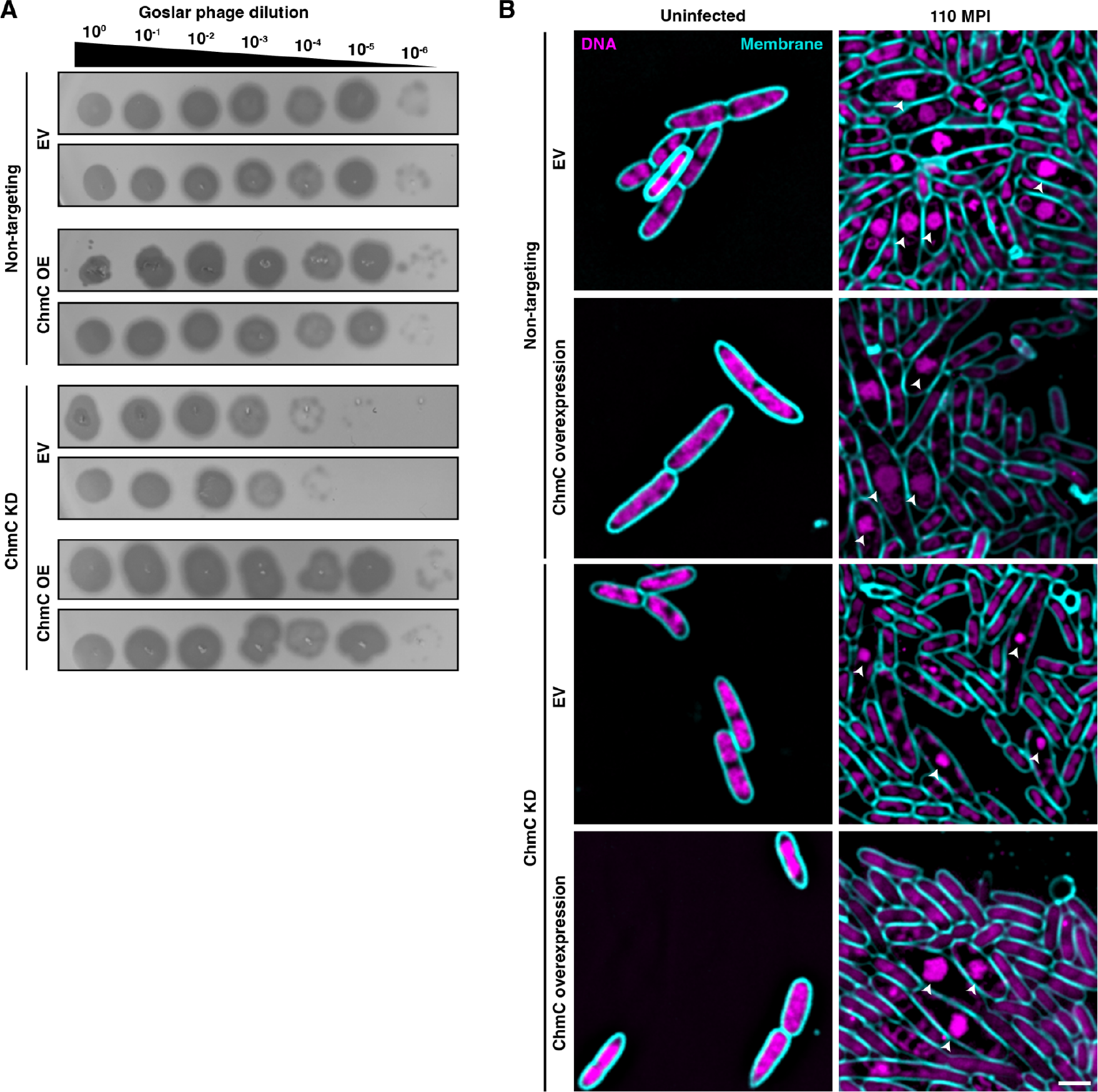
ChmC knockdown and complementation. **(A)** Plaque assays with phage Goslar on *E. coli* MC1000 cells expressing dCas13d and either a non-targeting guide RNA (NT) or a *chmC*-targeting guide RNA (ChmC KD). Host cells were additionally transformed with either an empty vector (EV) or a vector expressing recoded (dCas13d-resistant) ChmC (ChmC OE). See quantitation in **Figure 5B. (B)** Microscopy of *E. coli* MC1000 (uninfected at left; 110 MPI Goslar-infected at right) expressing dCas13d and either a non-targeting guide RNA (Non-targeting) or a *chmC*-targeting guide RNA (ChmC KD). Host cells were additionally transformed with either an empty vector (EV) or a vector expressing recoded (dCas13d-resistant) ChmC (ChmC overexpression). DNA is stained with DAPI and shown in magenta; membranes are stained with FM4-64 and shown in cyan. Arrowheads indicate the phage nucleus. Scale bar = 2 μm.

**Figure S7.**
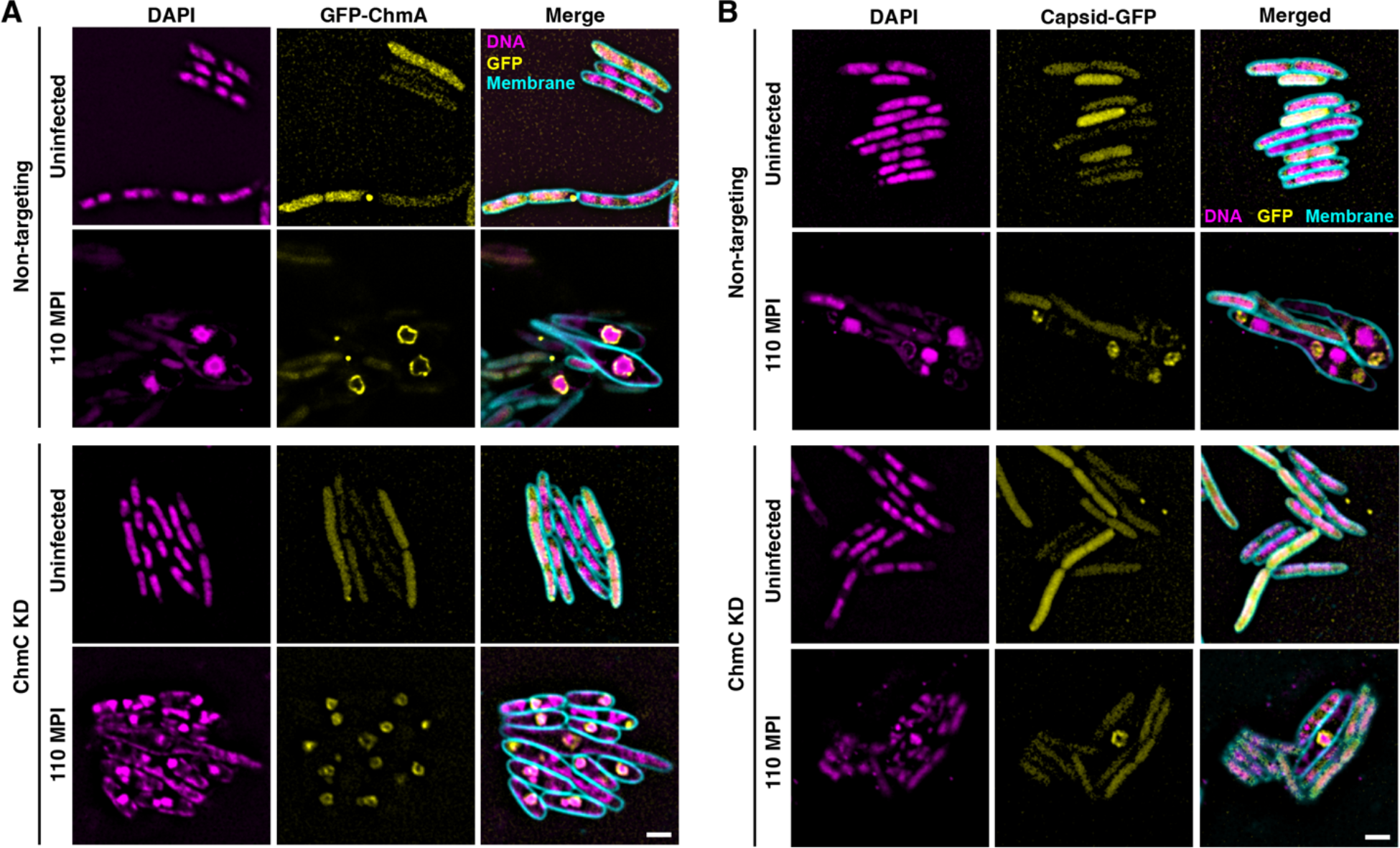
ChmC knockdown disrupts Goslar infections and causes a global protein knockdown. **(A)** Microscopy of *E. coli* MC1000 cells infected with Goslar (110 MPI) expressing GFP-tagged ChmA (gp246, yellow), dCas13d, and either a non-targeting guide RNA (top) or ChmC guide 3 (bottom). Arrowheads indicate phage nuclei, and asterisks indicate phage bouquets. **(B)** Microscopy of *E. coli* MC1000 cells infected with Goslar (110 MPI) expressing GFP-tagged phage capsid protein (gp41, yellow), dCas13d, and either a non-targeting guide RNA (top) or ChmC guide 3 (bottom). Arrowheads indicate phage nuclei, and asterisks indicate phage bouquets.

**Figure S8.**
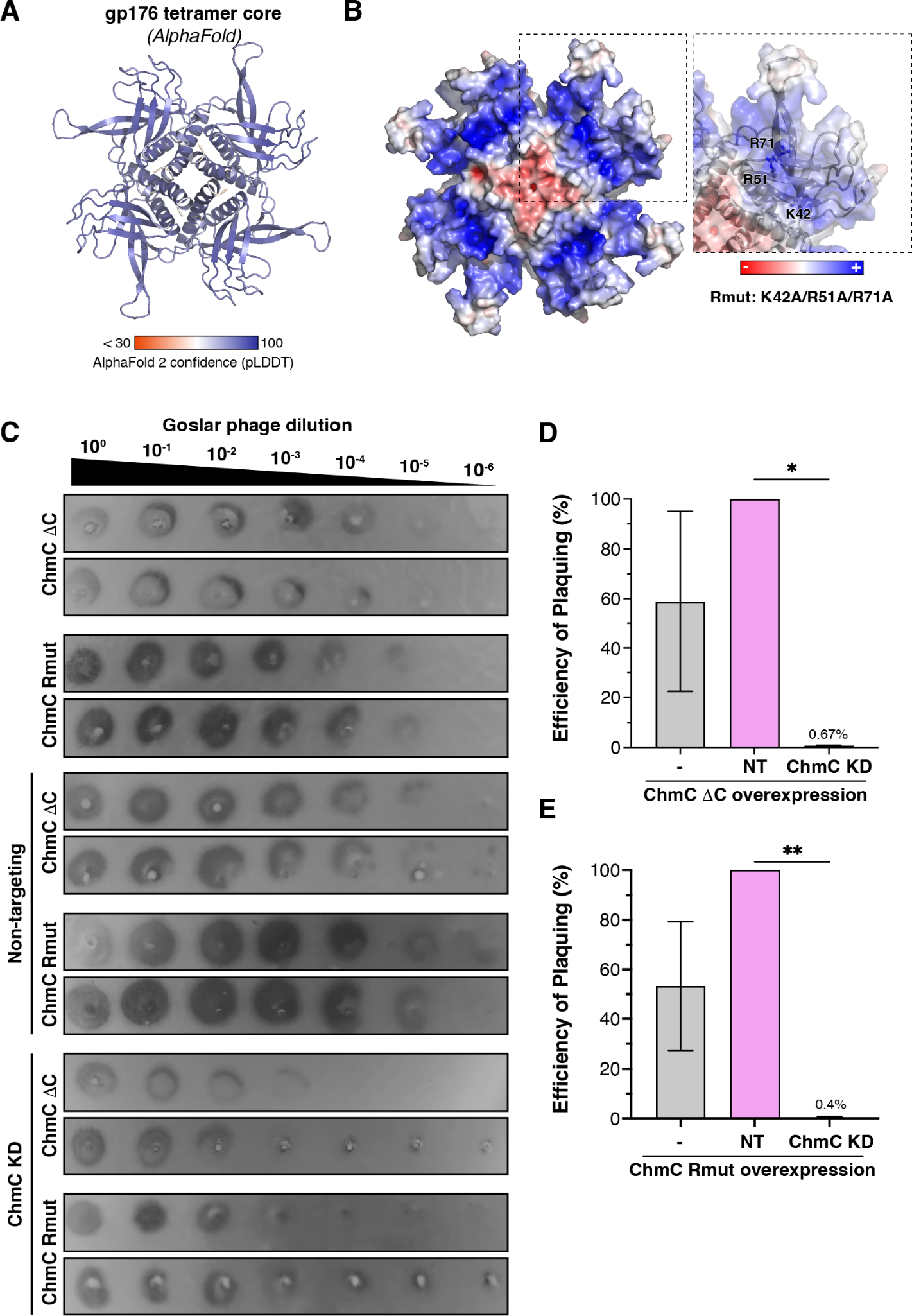
ChmC Rmut or ΔC overexpression does not rescue the knockdown phenotype. **(A)** AlphaFold predicted structure of a tetramer of Goslar ChmC (gp176), colored by prediction confidence (pLDDT score). Residues 16-192 of each chain are shown; N- and C-terminal tails with high predicted disorder are not shown. **(B)** View equivalent to panel (A) of a tetramer of Goslar ChmC (gp176), shown as a molecular surface colored by charge as calculated by APBS (46) in PyMOL. Closeup panel shows one monomer with residues equivalent to the PhiPA3 Rmut residues: K42, R51, and R71. **(C)** Plaque assays with phage Goslar on *E. coli* MC1000 cells overexpressing ChmC ΔC (residues 1-203) or Rmut (K42A/R51A/R71A), or expressing dCas13d and either a non-targeting guide RNA (NT) or a ChmC-targeting guide in addition to overexpressed ChmC ΔC or Rmut. **(D-E)** Efficiency of plaquing for the conditions shown in panel (C). * and ** indicate P values < 0.0332 and 0.0021 in a one-way ANOVA test of statistical significance.

## References

1. Hampton, H.G., Watson, B.N.J. and Fineran, P.C. (2020) The arms race between bacteria and their phage foes. Nature, 577, 327–336.

2. Stanley, S.Y. and Maxwell, K.L. (2018) Phage-Encoded Anti-CRISPR Defenses. Annu. Rev. Genet., 52, 445–464.

3. Prichard, A., Lee, J., Laughlin, T.G., Lee, A., Thomas, K.P., Sy, A., Spencer, T., Asavavimol, A., Cafferata, A., Cameron, M., et al. (2023) Identifying the core genome of the nucleus-forming bacteriophage family and characterization of Erwinia phage RAY. bioRxiv, 10.1101/2023.02.24.529968.

4. Chaikeeratisak, V., Nguyen, K., Khanna, K., Brilot, A.F., Erb, M.L., Coker, J.K.C., Vavilina, A., Newton, G.L., Buschauer, R., Pogliano, K., et al. (2017) Assembly of a nucleus-like structure during viral replication in bacteria. Science, 355, 194–197.

5. Chaikeeratisak, V., Nguyen, K., Egan, M.E., Erb, M.L., Vavilina, A. and Pogliano, J. (2017) The Phage Nucleus and Tubulin Spindle Are Conserved among Large Pseudomonas Phages. Cell Rep., 20, 1563–1571.

6. Knipe, D.M., Prichard, A., Sharma, S. and Pogliano, J. (2022) Replication Compartments of Eukaryotic and Bacterial DNA Viruses: Common Themes Between Different Domains of Host Cells. Annu Rev Virol, 9, 307–327.

7. Laughlin, T.G., Deep, A., Prichard, A.M., Seitz, C., Gu, Y., Enustun, E., Suslov, S., Khanna, K., Birkholz, E.A., Armbruster, E., et al. (2022) Architecture and self-assembly of the jumbo bacteriophage nuclear shell. Nature, 608, 429–435.

8. Mendoza, S.D., Nieweglowska, E.S., Govindarajan, S., Leon, L.M., Berry, J.D., Tiwari, A., Chaikeeratisak, V., Pogliano, J., Agard, D.A. and Bondy-Denomy, J. (2020) A bacteriophage nucleus-like compartment shields DNA from CRISPR nucleases. Nature, 577, 244–248.

9. Malone, L.M., Warring, S.L., Jackson, S.A., Warnecke, C., Gardner, P.P., Gumy, L.F. and Fineran, P.C. (2020) A jumbo phage that forms a nucleus-like structure evades CRISPR-Cas DNA targeting but is vulnerable to type III RNA-based immunity. Nat Microbiol, 5, 48–55.

10. Nieweglowska, E.S., Brilot, A.F., Méndez-Moran, M., Kokontis, C., Baek, M., Li, J., Cheng, Y., Baker, D., Bondy-Denomy, J. and Agard, D.A. (2023) The ϕPA3 phage nucleus is enclosed by a self-assembling 2D crystalline lattice. Nat. Commun., 14, 927.

11. Chaikeeratisak Vorrapon, Khanna Kanika, Nguyen Katrina T., Egan MacKennon E., Enustun Eray, Armbruster Emily, Lee Jina, Pogliano Kit, Villa Elizabeth and Pogliano Joe (2022) Subcellular organization of viral particles during maturation of nucleus-forming jumbo phage. Science Advances, 8, eabj9670.

12. Birkholz, E.A., Laughlin, T.G., Armbruster, E., Suslov, S., Lee, J., Wittmann, J., Corbett, K.D., Villa, E. and Pogliano, J. (2022) A cytoskeletal vortex drives phage nucleus rotation during jumbo phage replication in E. coli. Cell Rep., 40, 111179.

13. Enustun, E., Deep, A., Gu, Y., Nguyen, K.T., Chaikeeratisak, V., Armbruster, E., Ghassemian, M., Villa, E., Pogliano, J. and Corbett, K.D. (2023) Identification of the bacteriophage nucleus protein interaction network. bioRxiv, 10.1101/2023.05.18.541317.

14. Jumper, J., Evans, R., Pritzel, A., Green, T., Figurnov, M., Ronneberger, O., Tunyasuvunakool, K., Bates, R., Žídek, A., Potapenko, A., et al. (2021) Highly accurate protein structure prediction with AlphaFold. Nature, 596, 583–589.

15. Graebsch, A., Roche, S. and Niessing, D. (2009) X-ray structure of Pur-α reveals a Whirly-like fold and an unusual nucleic-acid binding surface. Proceedings of the National Academy of Sciences, 106, 18521–18526.

16. Schumacher, M.A., Karamooz, E., Zíková, A., Trantírek, L. and Lukes, J. (2006) Crystal structures of T. brucei MRP1/MRP2 guide-RNA binding complex reveal RNA matchmaking mechanism. Cell, 126, 701–711.

17. Desveaux, D., Allard, J., Brisson, N. and Sygusch, J. (2002) A new family of plant transcription factors displays a novel ssDNA-binding surface. Nat. Struct. Biol., 9, 512–517.

18. Mitrea, D.M., Cika, J.A., Guy, C.S., Ban, D., Banerjee, P.R., Stanley, C.B., Nourse, A., Deniz, A.A. and Kriwacki, R.W. (2016) Nucleophosmin integrates within the nucleolus via multi-modal interactions with proteins displaying R-rich linear motifs and rRNA. Elife, 5.

19. Pak, C.W., Kosno, M., Holehouse, A.S., Padrick, S.B., Mittal, A., Ali, R., Yunus, A.A., Liu, D.R., Pappu, R.V. and Rosen, M.K. (2016) Sequence Determinants of Intracellular Phase Separation by Complex Coacervation of a Disordered Protein. Mol. Cell, 63, 72–85.

20. Zaharias, S., Zhang, Z., Davis, K., Fargason, T., Cashman, D., Yu, T. and Zhang, J. (2021) Intrinsically disordered electronegative clusters improve stability and binding specificity of RNA-binding proteins. J. Biol. Chem., 297, 100945.

21. Azaldegui, C.A., Vecchiarelli, A.G. and Biteen, J.S. (2021) The emergence of phase separation as an organizing principle in bacteria. Biophys. J., 120, 1123–1138.

22. Nandana, V. and Schrader, J.M. (2021) Roles of liquid–liquid phase separation in bacterial RNA metabolism. Curr. Opin. Microbiol., 61, 91–98.

23. Lin, Y., Protter, D.S.W., Rosen, M.K. and Parker, R. (2015) Formation and Maturation of Phase-Separated Liquid Droplets by RNA-Binding Proteins. Mol. Cell, 60, 208–219.

24. Schmidt, H.B. and Görlich, D. (2016) Transport Selectivity of Nuclear Pores, Phase Separation, and Membraneless Organelles. Trends Biochem. Sci., 41, 46–61.

25. Chong, P.A., Vernon, R.M. and Forman-Kay, J.D. (2018) RGG/RG Motif Regions in RNA Binding and Phase Separation. J. Mol. Biol., 430, 4650–4665.

26. deYMartín Garrido, N., Orekhova, M., Lai Wan Loong, Y.T.E., Litvinova, A., Ramlaul, K., Artamonova, T., Melnikov, A.S., Serdobintsev, P., Aylett, C.H.S. and Yakunina, M. (2021) Structure of the bacteriophage PhiKZ non-virion RNA polymerase. Nucleic Acids Res., 49, 7732–7739.

27. Yakunina, M., Artamonova, T., Borukhov, S., Makarova, K.S., Severinov, K. and Minakhin, L. (2015) A non-canonical multisubunit RNA polymerase encoded by a giant bacteriophage. Nucleic Acids Res., 43, 10411–10420.

28. M Iyer, L., Anantharaman, V., Krishnan, A., Burroughs, A.M. and Aravind, L. (2021) Jumbo Phages: A Comparative Genomic Overview of Core Functions and Adaptions for Biological Conflicts. Viruses, 13.

29. Ceyssens, P.-J., Minakhin, L., Van den Bossche, A., Yakunina, M., Klimuk, E., Blasdel, B., De Smet, J., Noben, J.-P., Bläsi, U., Severinov, K., et al. (2014) Development of giant bacteriophage ϕKZ is independent of the host transcription apparatus. J. Virol., 88, 10501–10510.

30. Wicke, L., Ponath, F., Coppens, L., Gerovac, M., Lavigne, R. and Vogel, J. (2021) Introducing differential RNA-seq mapping to track the early infection phase for Pseudomonas phage ϕKZ. RNA Biol., 18, 1099–1110.

31. Armbruster, E.G., Lee, J., Hutchings, J., VanderWal, A.R., Enustun, E., Aindown Ann, A.B., Deep, A., Rodriguez, Z.K., Morgan, C.J., Ghassemian, M., et al. (2023) Sequential membrane- and proteinbound organelles compartmentalize genomes during phage infection. bioRxiv DOI: 10.1101/2023.09.20.558163.

32. Adler, B.A., Al-Shimary, M.J., Patel, J.R., Armbruster, E.G., Charles, E., Miller, K., Lahiri, A., Trinidad, M., Beurnier, S., Azadeh, A.L., et al. (2023) Targeted RNA knockdown unveils the functional genomes of bacteriophages. bioRxiv DOI: 10.1101/2023.09.18.558157.

33. Krupinska, K., Desel, C., Frank, S. and Hensel, G. (2022) WHIRLIES Are Multifunctional DNA-Binding Proteins With Impact on Plant Development and Stress Resistance. Front. Plant Sci., 13, 880423.

34. Luijsterburg, M.S., Noom, M.C., Wuite, G.J.L. and Dame, R.T. (2006) The architectural role of nucleoid-associated proteins in the organization of bacterial chromatin: a molecular perspective. J. Struct. Biol., 156, 262–272.

35. Dillon, S.C. and Dorman, C.J. (2010) Bacterial nucleoid-associated proteins, nucleoid structure and gene expression. Nat. Rev. Microbiol., 8, 185–195.

36. Hołówka, J. and Zakrzewska-Czerwińska, J. (2020) Nucleoid Associated Proteins: The Small Organizers That Help to Cope With Stress. Front. Microbiol., 11, 590.

37. Prikryl, J., Watkins, K.P., Friso, G., van Wijk, K.J. and Barkan, A. (2008) A member of the Whirly family is a multifunctional RNA- and DNA-binding protein that is essential for chloroplast biogenesis. Nucleic Acids Res., 36, 5152–5165.

38. Brocca, S., Grandori, R., Longhi, S. and Uversky, V. (2020) Liquid–Liquid Phase Separation by Intrinsically Disordered Protein Regions of Viruses: Roles in Viral Life Cycle and Control of Virus–Host Interactions. Int. J. Mol. Sci., 21, 9045.

39. Ye, Q., Lu, S. and Corbett, K.D. (2021) Structural Basis for SARS-CoV-2 Nucleocapsid Protein Recognition by Single-Domain Antibodies. Front. Immunol., 12, 719037.

40. Gales, J.P., Kubina, J., Geldreich, A. and Dimitrova, M. (2020) Strength in Diversity: Nuclear Export of Viral RNAs. Viruses, 12.

41. Richard Evans, Michael O’Neill, Alexander Pritzel, Natasha Antropova, Andrew Senior, Tim Green, Augustin Žídek, Russ Bates, Sam Blackwell, Jason Yim, et al. (2022) Protein complex prediction with AlphaFold-Multimer. bioRxiv.

42. Mirdita, M., Schütze, K., Moriwaki, Y., Heo, L., Ovchinnikov, S. and Steinegger, M. (2022) ColabFold: making protein folding accessible to all. Nat. Methods, 19, 679–682.

43. Blue, S.M., Yee, B.A., Pratt, G.A., Mueller, J.R., Park, S.S., Shishkin, A.A., Starner, A.C., Van Nostrand, E.L. and Yeo, G.W. (2022) Transcriptomewide identification of RNA-binding protein binding sites using seCLIP-seq. Nat. Protoc., 17, 1223–1265.

44. Lovci, M.T., Ghanem, D., Marr, H., Arnold, J., Gee, S., Parra, M., Liang, T.Y., Stark, T.J., Gehman, L.T., Hoon, S., et al. (2013) Rbfox proteins regulate alternative mRNA splicing through evolutionarily conserved RNA bridges. Nat. Struct. Mol. Biol., 20, 1434–1442.

45. Li, Q., Brown, J.B., Huang, H. and Bickel, P.J. (2011) Measuring reproducibility of high-throughput experiments. aoas, 5, 1752–1779.

46. Yee, B.A., Pratt, G.A., Graveley, B.R., Van Nostrand, E.L. and Yeo, G.W. (2019) RBP-Maps enables robust generation of splicing regulatory maps. RNA, 25, 193–204.

47. Schindelin, J., Arganda-Carreras, I., Frise, E., Kaynig, V., Longair, M., Pietzsch, T., Preibisch, S., Rueden, C., Saalfeld, S., Schmid, B., et al. (2012) Fiji: an open-source platform for biological-image analysis. Nat. Methods, 9, 676–682.

